# Morphospatial profiling of cancer-associated fibroblasts reveals architectural subtypes of pancreatic ductal adenocarcinoma

**DOI:** 10.64898/2026.02.06.704414

**Authors:** Adam S. Bryce, Silvia Martinelli, Leonor Schubert Santana, Leah Officer-Jones, Rachel L. Baird, Fiona Ballantyne, Kai Rakovic, Ian R. Powley, Dina Tataran, David J. Meltzer, Christopher Walsh, Fraser Duthie, Lucas Farndale, Adalberto Claudio Quiros, Ke Yuan, John Le Quesne, Fieke E.M. Froeling, Stephan B. Dreyer, David K. Chang

## Abstract

Pancreatic ductal adenocarcinoma (PDAC) is a lethal malignancy with an urgent need for biomarkers to predict prognosis and guide treatment. Understanding the complex spatial biology of pancreatic cancer-associated fibroblasts (CAFs) and the broader architecture of the PDAC tumour microenvironment is central to this challenge. Using a multi-omics approach across multiple spatial resolutions in a large human PDAC cohort, we integrate geometry and shape to define discrete morphological CAF subtypes, expanding CAF phenotyping beyond conventional proteomics. We then reveal an architectural and molecular axis of PDAC at tissue level, suggestive of epithelial-stromal co-evolution, with translational implications and prioritisation of stromal targets. Finally, we recapitulate this axis by introducing four unique, internally validated architectural subtypes of PDAC, each characterised by a common microenvironment and CAF enrichment profile. These archetypes outperform conventional pathology in prognostication, and predict response to adjuvant chemotherapy. Collectively, this study establishes a novel morphological paradigm for spatial biology, illuminates the architectural landscape of PDAC, and provides a framework for spatial biomarker discovery to close the translational gap in this devastating disease.

## Introduction

Pancreatic ductal adenocarcinoma (PDAC) is a lethal malignancy with dismal outcomes^1^. As current standard-of-care therapy offers only modest survival benefit, novel biomarkers to predict prognosis and guide treatment are urgently needed. A major challenge in this process is the complex PDAC tumour microenvironment (TME) and its constituent stroma, which represents up to 80% of tumour mass. Cancer-associated fibroblasts (CAFs) are key components of this stroma and are consistently implicated in tumour progression, microenvironmental crosstalk, and treatment response^2–6^.

Our understanding of CAF biology has increased in recent years, specifically CAF heterogeneity and functional diversity. However, previous work has predominantly focused on transcriptional states using bulk or single-cell techniques, and on pre-clinical models^7–14^. Key aspects of CAF biology remain elusive, including morphological heterogeneity, spatial organisation relative to other TME compartments, and contribution to global TME architecture^4^. Better understanding of these concepts will generate novel spatial biomarkers to accelerate therapeutic development and translation into the clinic^15,16^.

To improve our understanding of these morphospatial features, we performed multimodal molecular profiling on a large, clinically well-annotated cohort of patients with resected PDAC (n = 227) using multiplex immunofluorescence (mIF), genomics, and bulk transcriptomics. We then applied a hierarchical *pixel-to-patient* pipeline to achieve a comprehensive view of PDAC architecture across multiple spatial resolutions. At pixel level, we expanded CAF phenotyping beyond conventional spatial proteomics by integrating geometry and shape to define discrete morphological CAF subtypes. At cellular and neighbourhood levels, we profiled spatial interactions, stromal heterogeneity, and microenvironmental organisation to reveal an architectural axis of PDAC. Finally, at patient level, we introduced four unique and internally validated architectural subtypes (*archetypes*) of PDAC, each characterised by a common microenvironment. These archetypes were both prognostic and predictive, outperformed conventional histopathological variables, and were strongly linked to adjuvant chemotherapy response. Overall, this study establishes a framework for morphospatial biology, enabling discovery and development of biomarkers to predict prognosis and guide treatment in PDAC.

### Morphological and proteomic profiling defines biologically distinct CAF subtypes

Granular CAF characterisation was performed using low-plex mIF with key CAF markers on a cohort of treatment-naïve resected human PDAC tissue microarrays (TMAs) (n = 227, median three cores per patient) with detailed clinico-pathological annotations and mature follow-up (Fig. 1a, S1a-b). In addition, high-plex mIF with supplementary epithelial markers was performed on a subset of cases to profile the spatial relationships of epithelial subtypes of PDAC (n = 26). To investigate molecular associations, matching genomic (n = 182) and bulk transcriptomic (n = 59) datasets were also integrated (Fig. S1c). After staining and imaging, 2.5 million cells were segmented with a deep neural network and assigned to bulk cell types: CAF (FAP^POS^ CK^NEG^, 40%), Epithelial cell (FAP^NEG^ CK^POS^, 27%), Other cell (FAP^NEG^ CK^NEG^, 33%) (Fig. 1b-d). To faithfully capture their spindle shape, CAF boundaries were determined using a Laplacian transformation of FAP masks. Geometric features were then extracted and filtered to minimise inter-feature correlation, yielding a non-redundant set used to quantify CAF morphology at pixel resolution. This final set comprised five features: area, perimeter-to-area (PA) ratio, smoothness, elongation, and roundness (Fig. 1e, S1d).

**Fig. 1.**
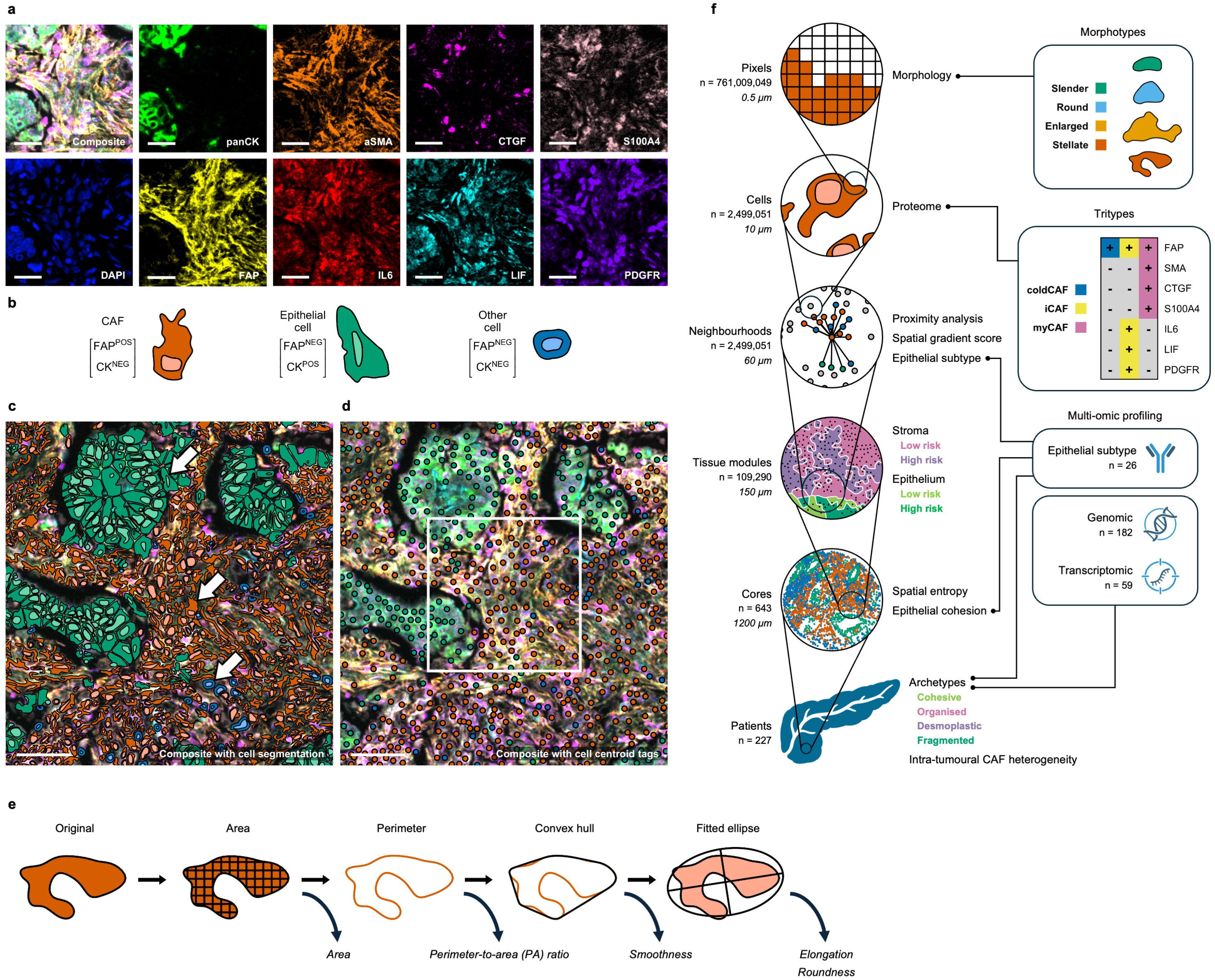
Pixel-to-patient analysis of PDAC spatial architecture and CAF morphology. **a,** Deep profiling of CAF biology and TME architecture was performed with a custom multiplex immunofluorescence panel designed and optimised for granular CAF characterisation. Scale bar = 30μm. **b,** 2.5 million cells were segmented with a deep neural network and initially assigned to bulk cell types. **c,** Full segmentation mask. To faithfully capture their spindle shape, CAF boundaries were determined using a Laplacian transformation of FAP masks. Scale bar = 50μm. **d,** Cell centroids only. Scale bar = 50μm. **e,** Computation of geometric features used to quantify CAF morphology at pixel resolution. **f,** Hierarchically profiling PDAC architecture with a *pixel-to-patient* pipeline across multiple spatial resolutions. Unit quantity and diameter (µm) (left) and computational techniques (right) are annotated. Epithelial subtypes and molecular patterns were determined with high-plex mIF (n = 26) and downstream integration of matched genomic (n = 182) and bulk transcriptomic (n = 59) datasets.

To achieve a comprehensive view of PDAC architecture, we designed a hierarchical *pixel-to-patient* analysis pipeline spanning pixel, cell, neighbourhood, tissue module, core, and patient resolutions (Fig. 1f). Distinct computational techniques at each level enabled extraction of biologically and clinically meaningful features, while preserving generalisability as features were aggregated.

Survival analyses revealed strong associations between geometric CAF features and clinical outcome, whereby roundness and smoothness were positively correlated with survival, while area and PA ratio were negatively correlated (Fig. 2a). Correlations between survival and protein marker intensities were comparatively modest, apart from CTGF and SMA expression which were negatively correlated (Fig. 2b). Noting globally low protein marker intensities in around 20% of CAFs, we then defined the “cold” pseudo-feature as the negative mean of all markers. This pseudo-feature positively correlated with survival and aligned with spindle-shaped geometry (roundness and smoothness). By contrast, CTGF and SMA expression strongly aligned with stellate-shaped features (area and PA ratio) (Fig. 2c).

**Fig. 2.**
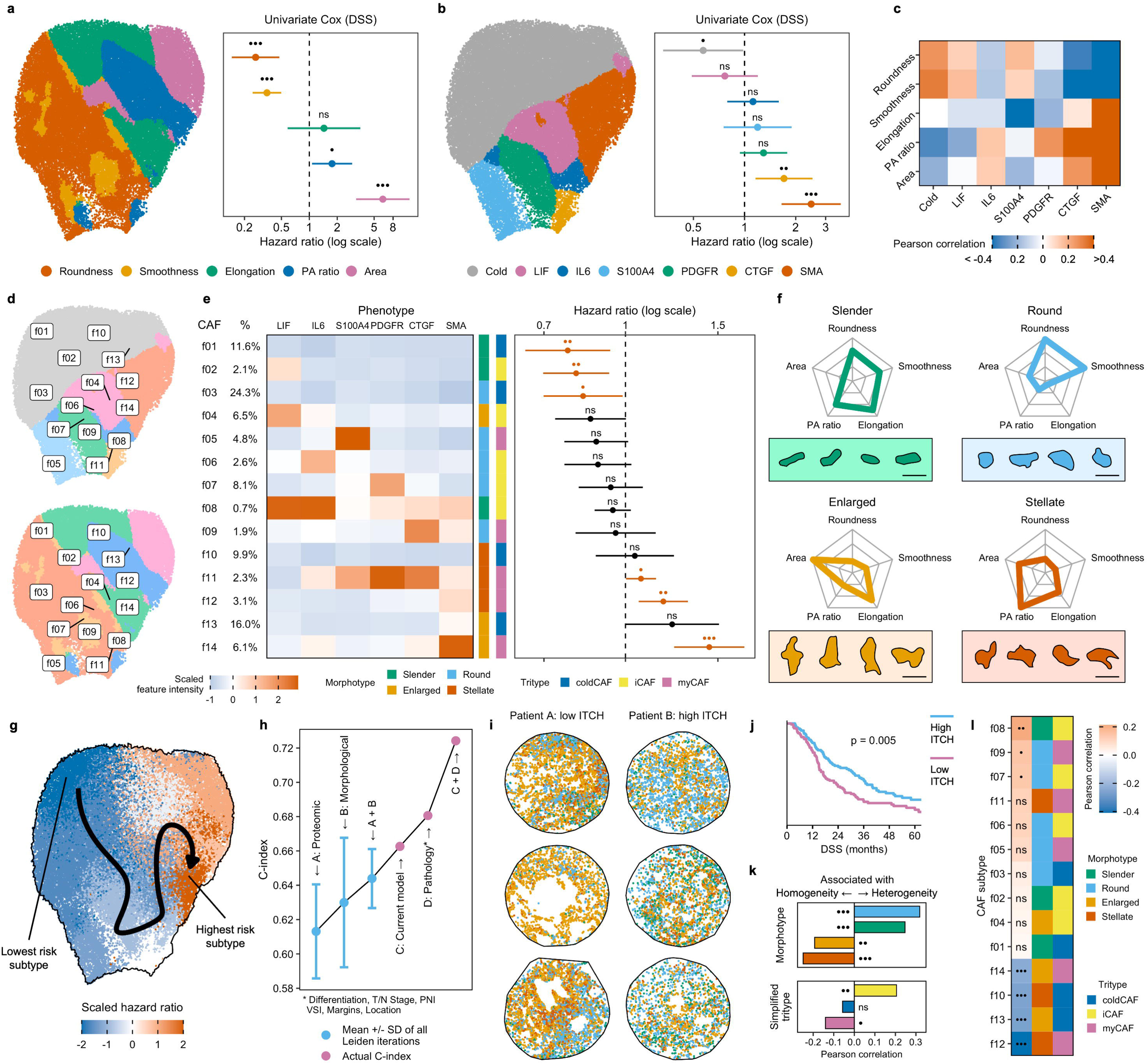
Morphological and proteomic profiling defines distinct CAF subtypes. **a-b,** Survival analyses revealed strong associations between CAF geometry and clinical outcome (**a**), while survival correlations with protein marker intensities were comparatively modest (**b**). DSS = disease-specific survival. UMAPs are false-coloured by neighbourhood maxima to convey local dominant feature. **c,** The “cold” pseudo-feature (the negative mean of all protein marker intensities) aligned with spindle-shaped geometry (roundness and smoothness), while CTGF and SMA strongly aligned with stellate-shaped features (area and perimeter-to-area ratio). **d-e,** Clustering of combined geometric and proteomic feature sets defined fourteen distinct CAF subtypes. Each subtype was then assigned to one of three *morphotypes* (geometric features) and one of three *tritypes* (proteomic features). CAFs broadly ranged from Slender and Round CAFs which were either proteomically cold or iCAF-like (neutral or favourable survival correlations), to Enlarged and Stellate CAFs with strong myCAF features (negative survival correlations). **f,** Feature enrichment and representative examples from each morphotype. Scale bar = 10µm. **g,** UMAP false-coloured by hazard ratio with line of best fit. **h,** Integrating geometric and proteomic feature sets yielded greater prognostic utility than each set in isolation. **i,** Quantification of intra-tumoural CAF heterogeneity (ITCH) with representative example cores (CAFs coloured by morphotype). **j,** High ITCH (stromal heterogeneity) was positively correlated with survival. p = Log-rank test (split by median). **k-l,** Correlations between ITCH and CAF enrichment by (**k**) morphotype and tritype and (**l**) all subtypes.

We next assigned CAF subtypes by combining geometric and proteomic feature sets with equal weighting. Using an iterative Leiden clustering pipeline, we identified fourteen distinct CAF subtypes, each with different survival associations (Fig. 2d-e, S2a-d). CAF subtypes were then assigned to one of four *morphotypes* (geometric features) and one of three *tritypes* (proteomic features). Morphotypes were named according to their dominant features – *Slender*, *Round*, *Enlarged*, or *Stellate*, while tritypes were named to align with established CAF classifications^4,7,8^ while integrating the cold pseudo-feature – *coldCAF*, *iCAF* (inflammatory CAF), or *myCAF* (myofibroblastic CAF) (Fig. 2f, S2e-h, S3a-b). This integrated approach, using both geometric and proteomic feature sets, demonstrated greater prognostic utility than each set in isolation (Fig. 2g-h).

These results demonstrate the utility of morphology in defining biologically and clinically meaningful CAF subtypes. We establish a relationship between CAF shape and proteome with prognostic significance, ranging from Slender and Round CAFs which are either proteomically cold or iCAF-like (neutral or favourable survival correlations), to Enlarged and Stellate CAFs with strong myCAF features (poor survival correlations). Furthermore, geometric features are applicable across cell types and disease contexts, and readily computable from cell boundary approximations alone. Morphological profiling therefore represents a transferable and accessible framework not only for CAF characterisation, but for spatial biology in general.

### CAF enrichment patterns in homogeneous and heterogeneous tumours

Intra-tumoural heterogeneity is a well-recognised feature of PDAC and is implicated in tumour evolution, therapy resistance, and metastasis^17–19^. However, recent assessment of heterogeneity has focused mostly on epithelial, immune, genomic, and transcriptomic signatures^20^. By focusing on stroma, we defined intra-tumoural CAF heterogeneity (ITCH) as the mean pairwise difference in CAF subtype enrichment across all TMA cores from each patient (Fig. 2i, S4). With a median of three spatially distinct cores per patient, ITCH aimed to capture intra-tumoural variation across regions, while using compositional normalisation to ensure fair comparison of cores and patients with varying stromal densities.

Initial analyses demonstrated better survival among patients with heterogeneous CAF enrichment (high ITCH) compared to those with homogeneous CAF enrichment (low ITCH) (Fig. 2j). Interrogating the contributions of specific CAF subtypes revealed correlations between high ITCH and the Slender and Round morphotypes, and the iCAF tritype. Conversely, correlations were noted between low ITCH and the Enlarged and Stellate morphotypes, and the myCAF tritype (Fig. 2k-l, S3c).

These results reveal a significant relationship between stromal heterogeneity, CAF subtypes, and survival. Tumours with a diverse, heterogeneous stroma were associated with improved survival and were enriched for small, round CAFs and iCAF-like states. Conversely, tumours with a uniform, homogeneous stroma were associated with poorer outcomes and were enriched for larger, stellate CAFs and myCAF-like states. This contrasts with the prevailing view of *epithelial* intra-tumoural heterogeneity in PDAC as a driver of tumour evolution, metastasis, and poor prognosis^20^. One possible explanation is a co-evolutionary process, whereby epithelial subclonal diversity expands and evolves under selective pressure^21,22^, while the stroma simultaneously loses diversity and converges towards a homogeneous, uniform state dominated by Enlarged and Stellate myCAF-like CAFs. Intra-tumoural CAF heterogeneity therefore provides a stromal-centric, biologically relevant metric and may act as a protective feature, representing tumours with high stromal diversity and more favourable outcomes.

### CAF subtypes localise to discrete spatial niches

After analysing CAF morphology, proteome, and heterogeneity with bulk tumour and single-cell approaches, we next examined the spatial relationships of CAFs with the wider TME, and their biological significance. This was performed by designating epithelial cells as the *anchor cells* from which the distances to each CAF were measured. To correct for anchor cell density, measurements were normalised by binning into *distance zones*, after which CAF subtype enrichment was computed within each zone in a custom tensor-based *proximity analysis* pipeline (Fig. 3a, S5).

**Fig. 3.**
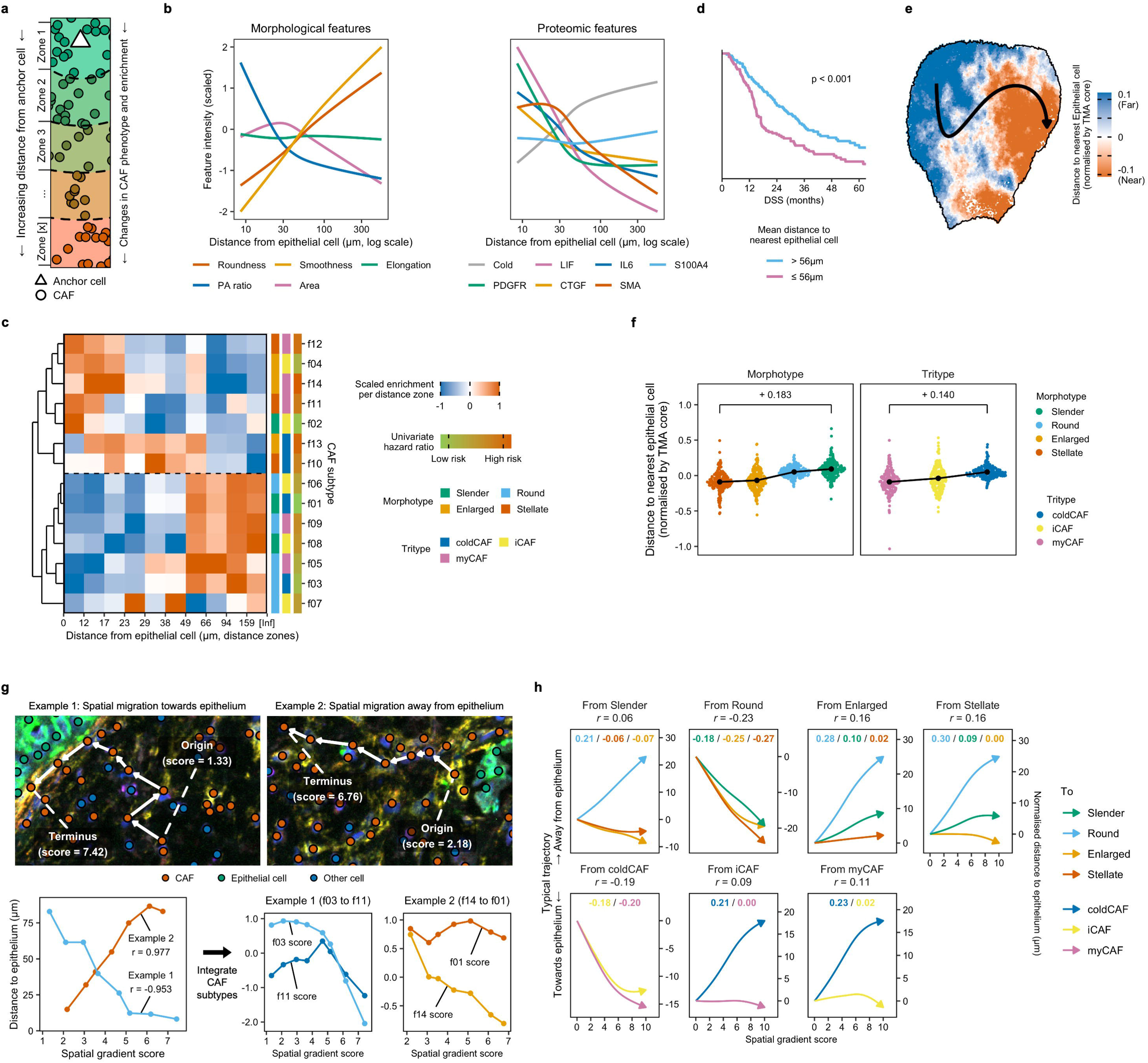
CAF subtypes localise to discrete spatial niches. **a,** *Anchor cell* and *distance zone* concepts. **b,** Relationships between raw morphological and proteomic features per distance zone. **c,** *Proximity analysis* revealed strong alignment of spatial niches with CAF morphotypes, and modest alignment with CAF tritypes. Distance zones are quantile defined. **d,** CAF separation from Epithelial cells correlated with survival. p = Log-rank test (split by median). **e,** UMAP false-coloured by normalised distance to nearest anchor cell with line of best fit. **f,** These findings were consistent across normalisation methods, suggesting they are robust to variations in anchor cell density. Stepwise changes between epithelium-adjacent and epithelium-distant subtypes are annotated, highlighting the strong alignment of spatial niches with CAF morphotypes, as compared to CAF tritypes. **g,** Overview of the *spatial gradient score*. Correlation between spatial gradient score and distance to epithelium was used to infer spatial trajectory towards epithelium (Example 1) or away from epithelium (Example 2) for each CAF subtype transition. **h,** Typical trajectories for each CAF subtype transition (aggregated by morphotype and tritype) with pairwise (inset) and global (title) Pearson correlation coefficients.

Exploratory analysis of raw morphological features revealed a marked relationship between CAF shape and spatial niche, whereby CAFs close to Epithelial cells were larger and more irregular in shape (high PA ratio) than those further away (Fig. 3b). Similar relationships were seen with proteomic features, most notably lower expression of SMA in CAFs further away from Epithelial cells. Expanding these findings to CAF subtypes using proximity analysis revealed consistent alignment of morphotypes and spatial niches (Fig. 3c). Enlarged and Stellate CAFs were enriched close to Epithelial cells, while Slender and Round CAFs were enriched further away. Although weaker, alignment was also noted with tritypes, whereby the myCAF tritype was enriched close to Epithelial cells and the iCAF tritype was enriched further away.

To ensure consistency of results while continuing to correct for anchor cell density, distances were then normalised per TMA core and subsequently aggregated by morphotype and tritype. This revealed a reproducible alignment of CAF subtypes and spatial niches, indicating that these relationships are robust to both inter-core variation in anchor cell density, and to normalisation method (Fig. 3e-f, S6a). In addition to biological relevance, CAF spatial relationships also held prognostic relevance, with separation of CAFs and epithelial cells correlating positively with survival (Fig. 3d). This correlation was also robust to variation in anchor cell density on survival modelling (Fig. S6b).

These results suggest that CAF subtypes are not randomly distributed, but consistently localise to discrete spatial niches, with the morphology and proteome of a CAF reflecting its local habitat. We hypothesise these dynamically influence each other, with crosstalk from neighbouring compartments providing molecular cues that shape CAF state and subtype. We further hypothesise that this could represent spatial plasticity, whereby CAFs reversibly transition between subtypes in response to local cues. Conversely, subtype transitions could drive migration of CAFs through the TME, redistributing them between spatial niches in alignment with changes in their subtype.

Our results also support a model of microenvironmental crosstalk in which CAFs actively influence epithelial biology in addition to passively responding to epithelial spatial cues. CAF-epithelial proximity was negatively correlated with survival, suggesting a hostile TME in which CAF-derived signals may be driving plasticity, heterogeneity, and metastatic potential in the neighbouring epithelium. Furthermore, the enrichment of Enlarged and Stellate CAFs in proximity to epithelium highlights these morphotypes as potential key drivers of this malignant progression.

### Spatial gradient scoring aligns CAF plasticity and epithelial proximity

To further investigate the relationship between CAF plasticity, spatial migration, and CAF-epithelial crosstalk, we developed a *spatial gradient score*. Hypothesising that spatially related CAFs may also be related in lineage, we aggregated neighbouring CAFs into paths using graph representation (Fig. S7). The gradient of subtype transition was used to assign scores along each path, and this was correlated with distance to epithelium. This integrative method, combining CAF plasticity (subtype transitions), spatial migration (paths), and microenvironmental crosstalk (epithelial distance), allowed subtype transitions to be characterised by their inferred spatial trajectories towards or away from epithelium (Fig. 3g).

Epithelial trajectories differed substantially across morphotype pairs, ranging from negative correlations, suggesting spatial migration towards epithelium (e.g. Round to Enlarged, *r* = - 0.25), to positive correlations, suggesting spatial migration away from epithelium (e.g. Enlarged to Slender, *r* = 0.10). When aggregated by morphotype of origin, the strongest correlations between spatial gradient score and epithelial distance were observed in paths arising from Round CAFs (*r* = −0.23) and Enlarged CAFs (*r* = 0.16). Additional heterogeneity was evident across tritype pairs, particularly in paths arising from the coldCAF tritype (*r* = - 0.19) and the myCAF tritype (*r* = 0.11) (Fig. 3h, S8).

These results demonstrate marked and heterogeneous relationships between spatial gradient score and epithelial distance, aligning CAF subtype transitions with spatial trajectories towards or away from epithelium. In doing so, our findings recapitulate subtype-specific spatial niches defined previously, while augmenting them with a spatially-linked migratory dimension. These results also further highlight the potential role of CAF-epithelial crosstalk as a driver of CAF plasticity and spatial migration, and define the CAF subtypes most strongly influenced by this crosstalk.

### Tissue modules distinguish low- and high-risk tumour architecture

After defining CAF subtypes and their spatial niches at pixel and cellular resolution, we then scaled the analysis to neighbourhood level to determine the nature and biological relevance of higher-order spatial interactions across the PDAC TME. Neighbourhoods were defined by clustering bulk cell type counts within a 30µm radius around each cell (Fig. S9). Hazard ratios and compartment assignments (stroma or epithelium) were then computed for each neighbourhood. This enabled stratification of neighbourhoods within each compartment into low- and high-risk, reflecting positive and negative correlations with survival, respectively (Fig. 4a).

**Fig. 4.**
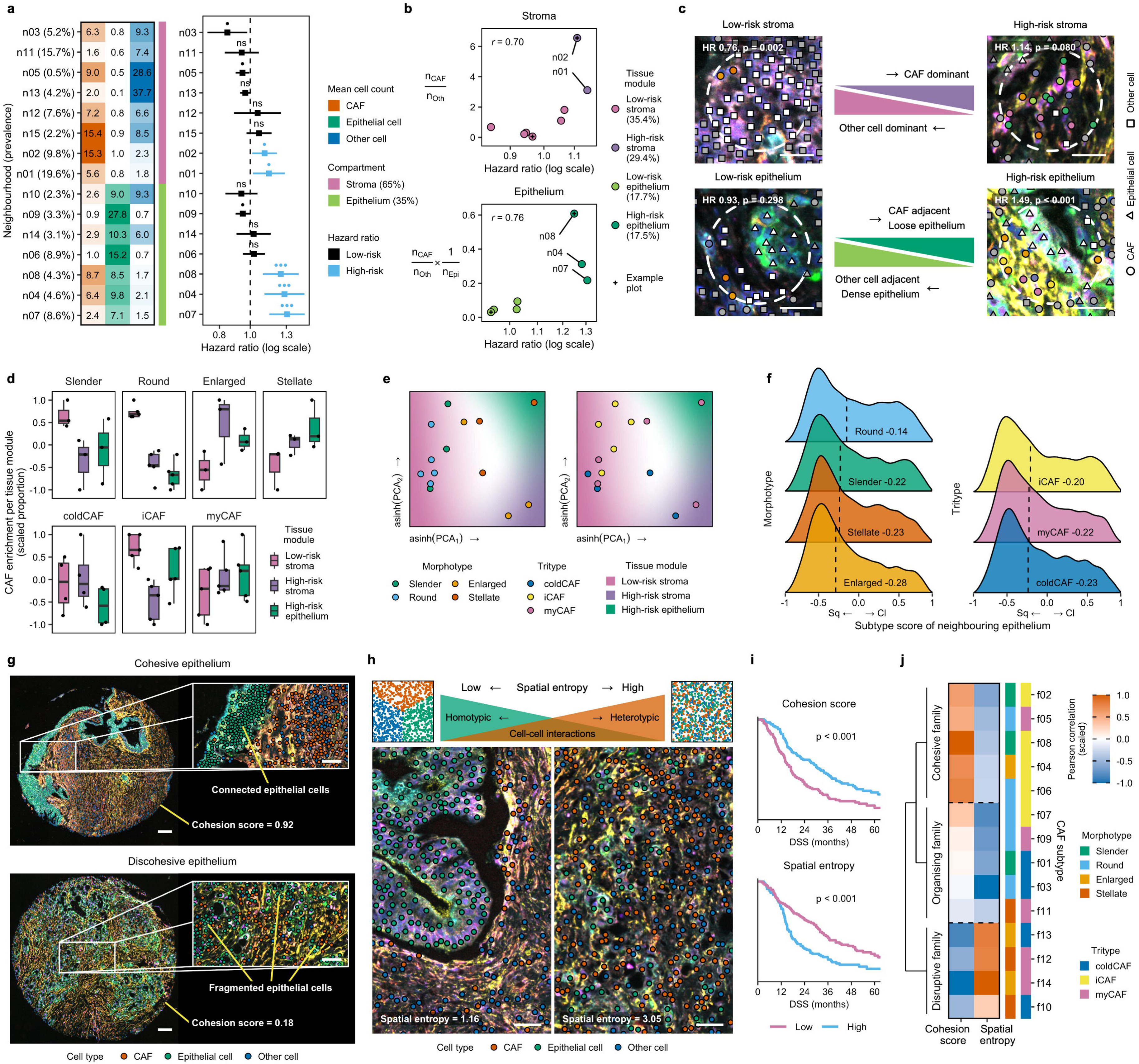
Tissue modules distinguish low- and high-risk tumour architecture. **a,** Overview of neighbourhoods, ordered by hazard ratio and compartment allocation (stroma or epithelium). **b,** Compartment-specific interaction terms (! axis) and neighbourhood hazard ratios (*x* axis) reduced fifteen neighbourhoods (n01 through n15) to four *tissue modules* (low-and high-risk stroma, and low- and high-risk epithelium). *r* = Pearson correlation. **c,** Examples of tissue modules from the extremes of each interaction term with survival correlations. Neighbourhood radius = 30µm. Scale bar = 20µm. **d-e,** CAF subtype enrichment per tissue module as boxplots (**d**) and PCAs (**e**). **f,** Heterogeneity of neighbouring epithelium by CAF subtype, ranging from 1 (most classical) to −1 (most squamous/basal-like). Annotation = median subtype score. **g-h,** *Cohesion score* and *spatial entropy* concepts. Scale bars = (**g**) 100µm/50µm, (**h**) 30µm **i,** High epithelial cohesion and low spatial entropy correlated with survival. p = Log-rank test (split by median). **j,** Correlating these features with CAF subtype enrichment revealed distinct architectural families for each morphotype and tritype.

To determine the differences in cellular composition between low- and high-risk neighbourhoods, we defined compartment-specific interaction terms. These revealed a negative correlation between survival and CAF count (both compartments), and positive correlations between survival and Other cell count (stroma), and the product of Other cell and Epithelial cell counts (epithelium) (Fig. 4b). The resulting alignment of interaction terms and hazard ratios reduced fifteen neighbourhoods to four *tissue modules*: low- and high-risk stroma, and low- and high-risk epithelium (Fig. 4c). In other words, neighbourhoods of comparable cellular composition aggregate into larger, distinct TME units, each representing the extremes of PDAC microenvironmental architecture with high prognostic relevance.

CAF morphotype composition varied widely across tissue modules, with enrichment of Slender and Round CAFs in low-risk stroma, Enlarged CAFs in high-risk stroma, and Stellate CAFs in high-risk epithelium (Fig. 4d-e). Although CAF tritype composition was more conserved in comparison, there was notable enrichment of the iCAF tritype in low-risk stroma. CAF subtype enrichment in low-risk epithelium could not be computed as this tissue module was entirely CAF-excluded.

These results reveal the marked architectural heterogeneity of the PDAC TME. Differences in cellular composition between low- and high-risk tissue modules suggest survival is influenced not by bulk cell density alone, but by the cumulative effect of a complex network of local cellular interactions and crosstalk throughout the TME. In particular, the contrast between low- and high-risk epithelium underscores the biological significance of dense, CAF-excluded, and loose, CAF-adjacent epithelial architecture, respectively. We also observed wide variation in CAF subtype enrichment between tissue modules which recapitulated subtype-specific spatial niches defined earlier. In particular, our findings further support Enlarged and Stellate CAFs as potential drivers of poor outcome in stromal and epithelial compartments, respectively. Notably, the association of Stellate CAFs with loose, fragmented epithelium implicates possible crosstalk between these cells as a key contributor to adverse epithelial biology.

### Neighbourhood heterogeneity in classical and squamous/basal-like epithelium

To investigate the relationship between CAF biology and epithelial subtypes in PDAC, we profiled a subset of patients (n = 26) with a high-plex custom mIF panel comprising canonical squamous/basal-like and classical markers^23–28^. To fully capture epithelial heterogeneity, each epithelial cell was assigned a continuous subtype score ranging from −1.0 (most squamous/basal-like) to +1.0 (most classical) (Fig. S10a-b). Higher mean subtype scores (i.e. more classical) were positively correlated with survival, and subtype scores corresponded to previously defined bulk transcriptomic subtypes, supporting the validity and biological relevance of this approach (Fig. S10c-d).

To determine the spatial associations between epithelial and CAF subtypes we applied previously defined 30µm windows to compute the mean subtype score of all epithelial cells within each local CAF neighbourhood. This revealed distinct CAF morphotype associations, most notably Round CAFs with classical-leaning and Enlarged CAFs with squamous/basal-like-leaning epithelium (median neighbourhood subtype scores −0.14 and −0.28, respectively). Modest correlations were also observed with CAF tritypes (Fig. 4f).

While local epithelial habitat is therefore reflected in CAF identity, we noted an overarching trend towards squamous/basal-like neighbourhoods across all CAF subtypes. To further investigate this, we reorientated our analysis to epithelial cell neighbourhoods and quantified their cellular compositions. This revealed that classical epithelial cells were more frequently found with no neighbouring CAFs, whereas squamous/basal-like epithelial cells were more frequently isolated from neighbouring epithelial cells (Fig. S10e).

These results demonstrate a clear relationship between epithelial subtype heterogeneity and local neighbourhood context. The association of squamous/basal-like cells with CAF-dense, epithelium-isolated neighbourhoods further implicates this architectural phenotype – closely resembling the previously defined high-risk epithelium tissue module – as a hallmark of poor tumour biology. Conversely, the association of classical cells with CAF-excluded, epithelium rich neighbourhoods recapitulates the previously defined low-risk epithelium tissue module. Together with the distinct spatial correlations observed between epithelial subtype and CAF morphotypes, these findings point to a complex interplay between CAF plasticity, epithelial plasticity, and microenvironmental architecture.

### Quantifying global differences in TME architecture

Having established the biological significance of epithelial organisation with neighbourhood and tissue module approaches, we next investigated the architectural heterogeneity of the broader PDAC TME by further scaling our analysis to whole TMA core level. To formally quantify epithelial organisation, a density-based *cohesion score* was developed to provide a continuous measure of epithelial connectivity, ranging from dense, highly connected epithelium (cohesion score →1), to loose, fragmented epithelium (cohesion score →0) (Fig. 4g). In parallel, we applied *spatial entropy*, a previously defined spatial form of Shannon entropy^29^, to capture TME-wide architectural organisation ranging from low (organised) to high (disorganised) (Fig. 4h).

Each architectural metric proved to be highly biologically informative, with strong survival associations observed for high cohesion score and low spatial entropy (Fig. 4i). Surprisingly, both were minimally correlated, suggesting they captured independent facets of tumour architecture (Fig. S11a). Given their apparent visual similarity to tumour differentiation, we then assessed their correlations with histological grade. Cohesion score showed a clear association, whereas spatial entropy did not, suggesting spatial entropy may capture architectural disorganisation not readily identified by routine histological assessment alone (Fig. S11b). Cohesion score also correlated strongly with epithelial subtype score, with cohesive tumours trending towards classical epithelial subtype and fragmented tumours towards squamous/basal-like epithelial subtype (Fig. S11c-d).

We also observed marked heterogeneity in CAF subtype enrichment across both metrics. This was most evident between Round and Enlarged CAFs and spatial entropy (*r* = −0.55 and *r* = 0.52, respectively), and between the iCAF and myCAF tritypes and cohesion score (*r* = 0.32 and *r* = −0.22, respectively) (Fig. S11e-f). Subsequent clustering of these enrichment patterns revealed three distinct architectural families, each with characteristic associations: *Cohesive family* (predominantly the iCAF tritype) – high epithelial cohesion, *Organising family* (predominantly Round and Slender CAFs) – low spatial entropy, and *Disruptive family* (predominantly Enlarged and Stellate CAFs, and the myCAF tritype) – low epithelial cohesion and high spatial entropy (Fig. 4j).

These results recapitulate architectural patterns defined at neighbourhood and tissue level and generalise them to the broader TME. We demonstrate the prognostic significance of a spatially organised TME with preserved architecture, connected epithelium, and enrichment of Cohesive and Organising CAFs within a contained stroma. Conversely, a disorganised TME with loss of compartmentalisation, enrichment of Disruptive CAFs, and abundant CAF-epithelial interactions was significantly associated with poor outcome. We hypothesise that such epithelial fragmentation reflects epithelial-mesenchymal transition (EMT), marking a critical breakdown of tumour architecture, acquisition of squamous/basal-like properties, and the emergence of metastatic potential.

### Architectural subtypes of PDAC

To explore the relationship between these architectural features and key clinical outcomes including survival and treatment response, we derived a final, high-order summary of tumour architecture at patient level by computing the dominant tissue module (Fig. 5a). Here, we introduce four architectural subtypes (*archetypes*) of PDAC, each characterised by a common microenvironment and specific CAF subtype enrichment. These archetypes were strongly prognostic and were defined as follows (dominant tissue module, median disease-specific survival): *Cohesive* (low-risk epithelium, 40.0 months), *Organised* (low-risk stroma, 31.0 months), *Desmoplastic* (high-risk stroma, 17.4 months), *Fragmented* (high-risk epithelium, 13.2 months) (Fig. 5b-c).

**Fig. 5.**
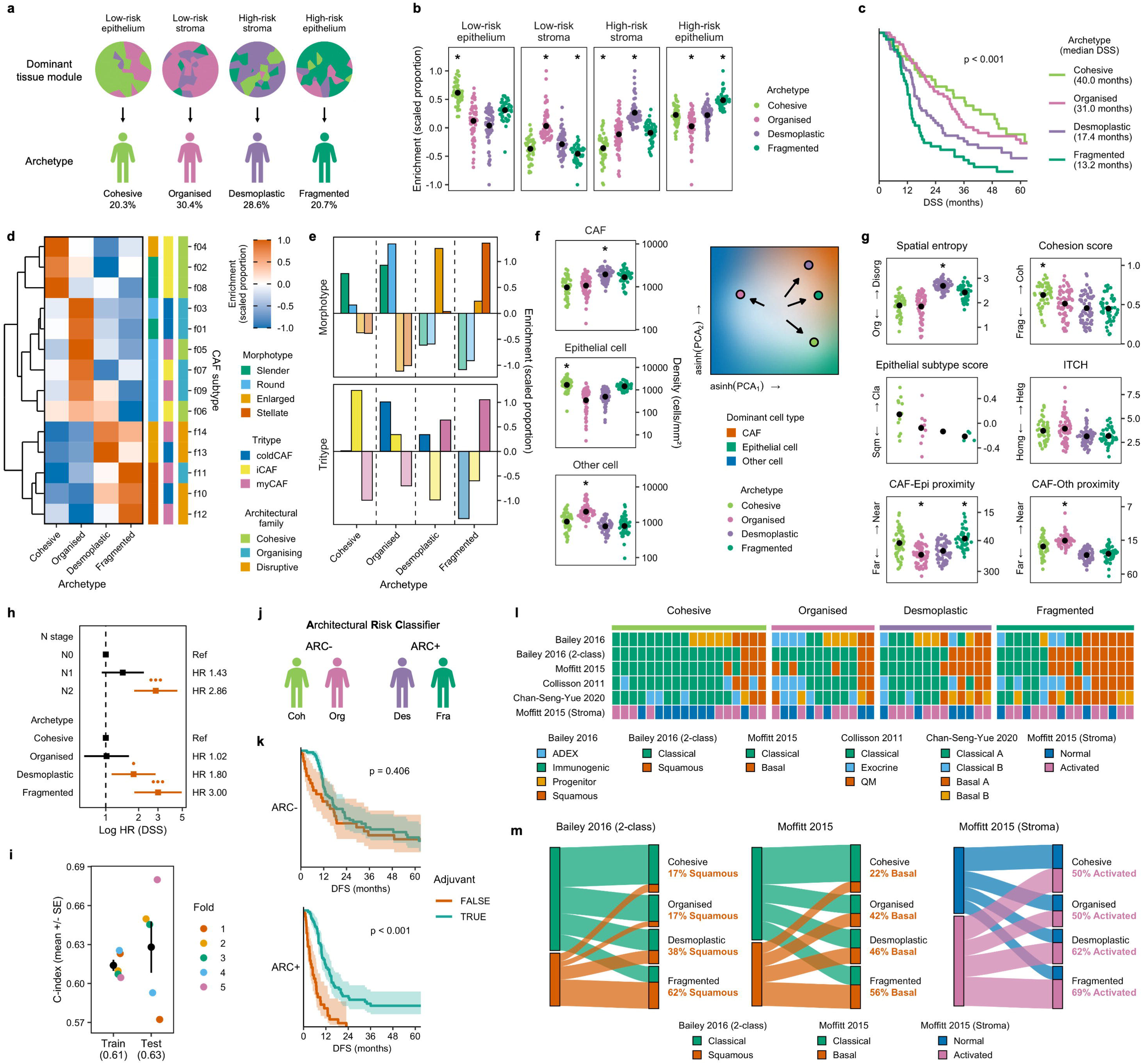
Architectural subtypes of PDAC. **a,** Dominant tissue module defines architectural subtype (*archetype*). **b,** Each archetype was characterised by a common microenvironment. Point = mean. * = key feature (minimum/maximum group AND four significant groupwise comparisons by T-test, applies to entire figure). **c,** Archetypes correlate strongly with survival. p = Log-rank test. **d-f,** Each archetype demonstrated distinct CAF morphotype and tritype enrichment (**d-e**), and bulk cell type profile (**f**). **g,** Archetypes were further characterised by broader microenvironmental differences. **h,** Archetypes outperformed conventional histopathological features on multivariate Cox regression (final model after backwards elimination with significance threshold of 0.01). **i,** The stability of archetype prognostication was confirmed with internal validation. **j-k,** Architectural Risk Classifier (ARC) predicted post-adjuvant disease-free survival (DFS), driven by significantly lower DFS in ARC+ patients who did not receive adjuvant treatment. p = Log-rank test. **l-m.** Archetypes align with molecular spectra of PDAC, particularly the classical to squamous/basal-like axis.

Each archetype demonstrated distinct enrichment of CAF morphotypes and tritypes (Fig. 5d-e). The Cohesive archetype was enriched for Slender iCAFs, and the Organised archetype for Round CAFs. myCAF enrichment was most pronounced in the Desmoplastic and Fragmented archetypes, with Enlarged myCAFs dominating in the former, and Stellate myCAFs in the latter. When aggregated by CAF architectural family, we observed clear alignment of Cohesive and Organising CAFs with Cohesive and Organised archetypes, and of Disruptive CAFs with Desmoplastic and Fragmented archetypes. Bulk cell type profiles also varied, with enrichment of Epithelial cells in the Cohesive archetype, and of CAFs in the Desmoplastic archetype (Fig. 5f). Archetypes were further characterised by broader microenvironmental differences: Cohesive – high cohesion score, Organised – low spatial entropy, Desmoplastic – low ITCH, Fragmented – low cohesion score and high CAF-Epithelial cell proximity (Fig. 5g). We also noted a clear shift towards squamous/basal-like epithelial subtype scores in Desmoplastic and Fragmented archetypes.

Multivariate Cox regression analysis confirmed the independent prognostic value of architectural subtyping beyond conventional histopathological features (Fig. 5h, S12a). Additionally, minimal variation was observed between archetypes across demographics, histopathological features, and recurrence pattern (Fig. S12b). We also confirmed the stability and generalisability of archetype prognostication by internally validating with a five-fold cross-validation technique, which yielded concordance indices (mean ± SD) of 0.61 ± 0.01 in training and 0.63 ± 0.04 in testing (Fig. 5i, S13).

Building on this prognostic utility, we evaluated the predictive value of architectural subtyping in adjuvant chemotherapy response by examining disease-free survival (DFS). We defined an Architectural Risk Classifier (ARC) by aggregating Cohesive and Organised archetypes to ARC-, and Desmoplastic and Fragmented archetypes to ARC+, representing low- and high-risk tissue modules, respectively (Fig. 5j, S14a-c). This classifier demonstrated marked predictive utility (adjuvant × ARC interaction HR = 0.32, 95% CI 0.16-0.61, p=0.002), reflecting a significant decrease in DFS in ARC+ patients who did not receive adjuvant treatment (13.0 vs 4.9 months, p<0.001), and a non-significant decrease in similar ARC- patients (22.0 vs 17.3 months, p=0.406) (Fig. 5k, S14d-f). The stability and generalisability of this classifier were confirmed by internal cross-validation, which yielded adjuvant × ARC interaction HRs (mean ± SD) of 0.33 ± 0.07 in training and 0.51 ± 0.46 in testing (Fig. S14g-h).

### Architectural subtypes map to molecular programmes

To uncover the molecular programmes underlying these architectural subtypes, we integrated matching genomic (n = 182) and bulk transcriptomic (n = 59) datasets. Analysis of canonical PDAC driver genes (*KRAS*, *TP53*, *SMAD4*, and *CDKN2A*) revealed conserved mutational profiles and consistent *KRAS* variants across archetypes, suggesting that architectural heterogeneity arises downstream of common genomic events (Fig. S15a-c). In contrast, gene set enrichment analysis of bulk transcriptomes using the MSigDB Hallmarks library revealed archetype-specific molecular programmes (Fig. S15d-f). Immune and metabolic pathways, such as interferon response and oxidative phosphorylation, were upregulated in the Cohesive archetype, while stromal-associated programmes, such as myogenesis, were downregulated in the Cohesive compared to the Desmoplastic archetype. Analysis using the IOBR reference library^30^ revealed similar upregulation of immune and metabolic signatures in the Cohesive archetype, particularly T-cell activation, antigen presentation, and lipid metabolism pathways.

Archetypes were then mapped to previously described bulk transcriptomic PDAC subtypes to further characterise their molecular profiles^23,31–33^ (Fig. 5l-m, S15g). Significant differences were observed for the Fragmented archetype, with enrichment of the Bailey squamous and Collisson quasi-mesenchymal subtypes, and depletion of the Collisson classical subtype. A similar trend from classical to squamous/basal-like status across the archetype spectrum was evident when applying Moffitt and Chan-Seng-Yue classifications. Further examination revealed enrichment of the Bailey immunogenic subtype in the Cohesive archetype, and depletion of the Bailey progenitor subtype in the Fragmented archetype. Finally, application of the Moffitt stromal classification revealed a parallel trend from normal to activated stroma across the archetype spectrum.

These results demonstrate the utility of architectural subtyping in identifying biologically and clinically distinct variants of PDAC, each with characteristic spatial features and CAF enrichment signatures (Fig. 6a-d). Weak correlations between archetypes and histopathological features also suggest that architectural subtyping is capturing a unique axis of tumour biology not represented in conventional histological assessment. Furthermore, the strong interaction between adjuvant treatment and ARC status suggests that adjuvant response is augmented in spatially disorganised, CAF-dense tumours with fragmented epithelium. This finding may reflect a higher risk of recurrence among patients with more aggressive disease biology, thereby identifying those more likely to benefit from adjuvant chemotherapy.

**Fig. 6.**
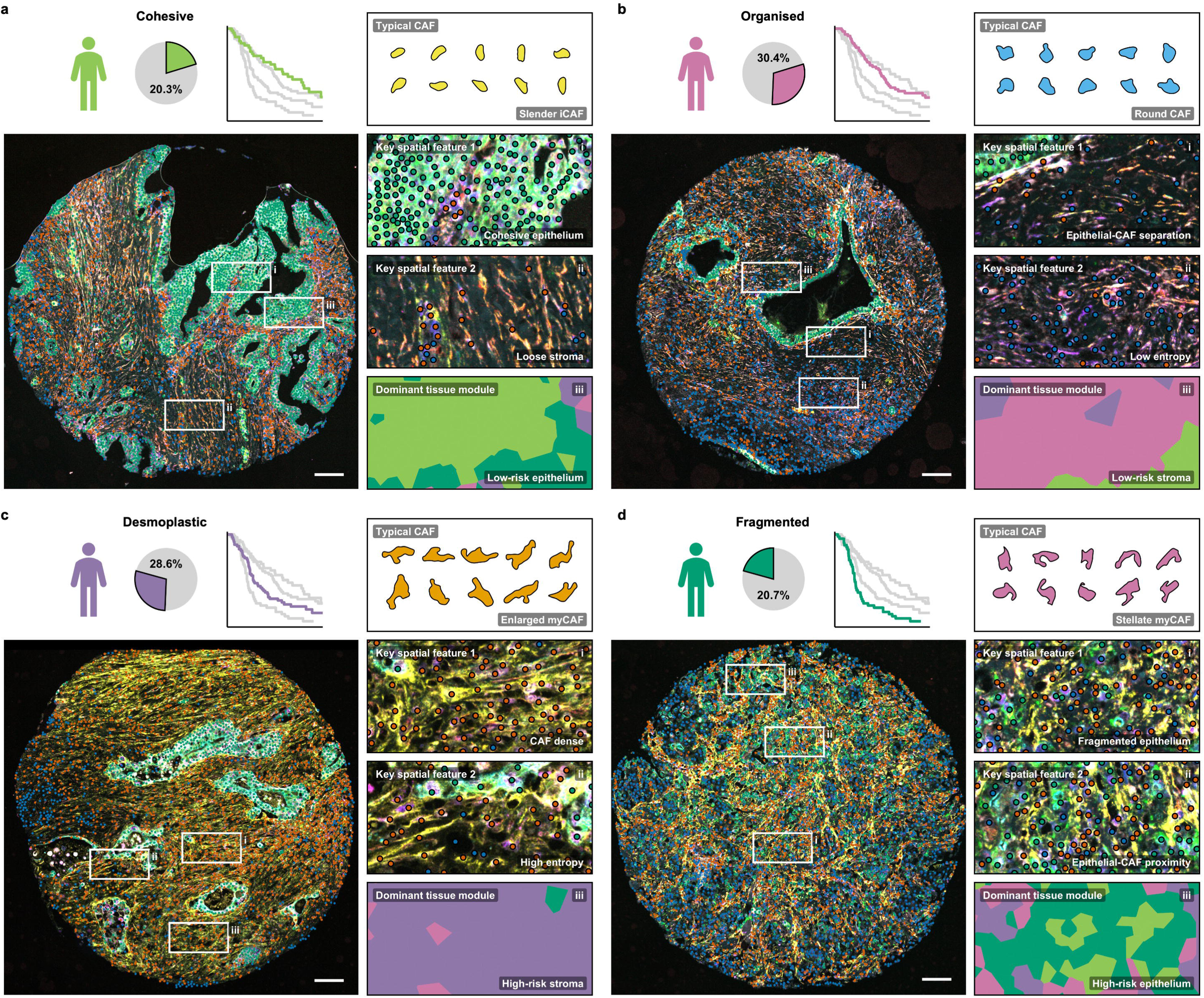
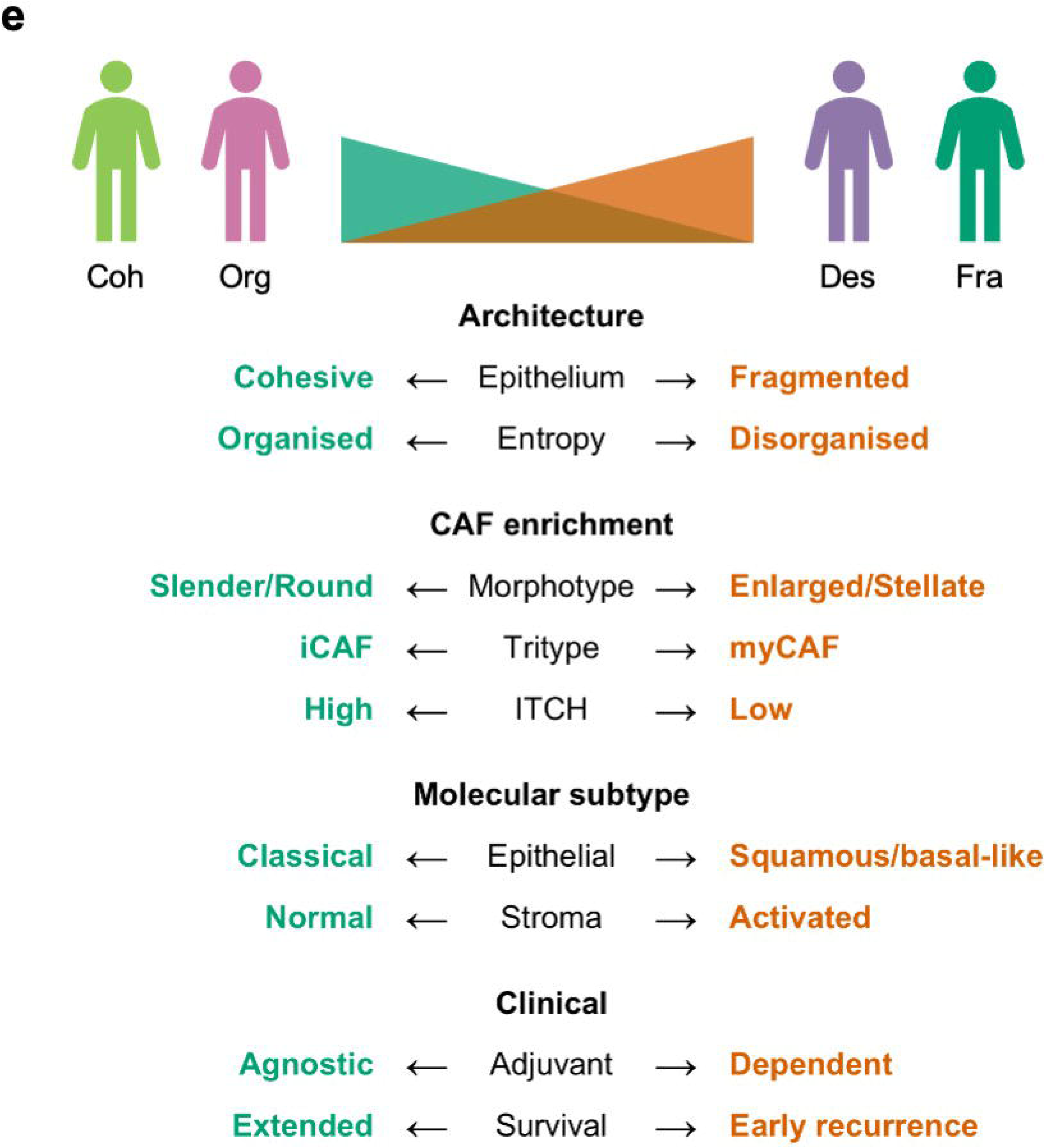
Overview of archetypes and co-evolutionary model. **a-d,** Example core, typical CAF, key features, and dominant tissue module of each archetype. Scale bar = 100µm. **e,** A unifying co-evolutionary model of PDAC across the archetype spectrum, defined by parallel architectural, stromal, molecular, and clinical axes.

These results also align PDAC architectural heterogeneity to distinct molecular mechanisms. The transcriptional signatures of the Cohesive archetype indicate an immune-competent, T-cell infiltrated microenvironment mapped to the Bailey immunogenic subtype, with corresponding spatial enrichment of Slender iCAFs. A classical-leaning transcriptome and prominent metabolic programmes also suggest that the epithelial cohesion which defines this archetype is recapitulating ductal architecture and pancreatic lineage. Conversely, transcriptional patterns in the Fragmented archetype support a dedifferentiated, progenitor depleted, squamous/basal-like lineage, corresponding to epithelial fragmentation, architectural breakdown, and high metastatic potential. Furthermore, enrichment of myogenesis signatures from the Cohesive to the Desmoplastic archetype reflects the compositional change from Slender and Round CAFs to Enlarged and Stellate myCAFs across the archetype spectrum. This suggests an axis of stromal organisation, with a parallel increase in CAF-Epithelial cell proximity, loss of intra-tumoural CAF heterogeneity, and a transcriptomic shift from Moffitt normal to activated stromal signatures (Fig. 6e).

## Discussion

Pancreatic ductal adenocarcinoma is a deadly malignancy and remains a cancer of unmet need, with little improvement in survival over recent decades. In this study, we sought to characterise the PDAC tumour microenvironment, particularly its complex stroma, by profiling CAFs and broader tumour architecture in a large, well characterised patient cohort using a hierarchical pixel-to-patient approach spanning multiple spatial resolutions. By understanding these processes, we aim to identify clinically relevant spatial biomarkers to enable stratification of treatment and further discovery of therapeutic targets.

Although our understanding of CAF heterogeneity in PDAC has increased exponentially in recent years, most classification frameworks are derived from transcriptomic and proteomic data^34^, with little focus on cell shape. In this study, we reveal that CAF morphology encodes rich biological information, outperforms low-plex proteomics in CAF phenotyping, and aligns faithfully with epithelial proximity and microenvironmental niche. Furthermore, morphology can be determined from single-plex cell boundary approximations, making it an accessible, cost-effective, and disease-agnostic tool that is readily compatible with current routine clinical, histopathological, and spatial proteomic workflows.

Spatial analysis of morphotypes revealed enrichment of Enlarged and Stellate CAFs in peri-epithelial zones. Consistent with prior reports, myCAFs were similarly concentrated in these zones^7,35,36^. Poorer outcomes were also observed in tumours with higher peri-epithelial CAF density. Aligning with previously published work demonstrating a spindle-to-stellate transition during CAF activation^37^, these results suggest that Enlarged and Stellate myCAFs represent a highly activated CAF state. Proximity to epithelium and association with poor survival imply these CAF subtypes may also be engaged in reciprocal crosstalk that could catalyse malignant behaviour in the neighbouring epithelium, highlighting these cells as potential stromal drivers of poor outcome.

Scaling our analysis to neighbourhoods and tissue modules, we demonstrate a continuum of CAF enrichment across archetypes, from Slender iCAFs and Round CAFs in Cohesive and Organised archetypes, to Enlarged and Stellate myCAFs in Desmoplastic and Fragmented archetypes. Profiling of epithelial cohesion, spatial entropy, intra-tumoural CAF heterogeneity, and epithelial/stromal subtype revealed parallel axes of microenvironmental organisation, stromal composition, and molecular programmes across the archetype spectrum (Fig. 6e). Together, these findings could support a co-evolutionary model in which the epithelium shifts towards a fragmented, disorganised, squamous/basal-like phenotype, while the stroma converges on a homogeneous, transcriptomically activated state enriched for Enlarged and Stellate myCAFs. This concept aligns with prior reports linking epithelial to stromal co-evolution^38,39^, and myCAF spatial abundance to squamous/basal-like epithelium^35,40^.

Therapeutically, this model poses two questions of significant clinical relevance. First, are these stromal and epithelial shifts irreversible, or do they retain plasticity that could be leveraged to stabilise tumour architecture? Second, given their co-evolutionary coupling, would arresting stromal progression in turn halt epithelial evolution? To guide this hypothesis, we highlight potential stromal targets across spatial resolutions; at cell level, targeting high risk CAF states (Enlarged and Stellate myCAFs), at neighbourhood level, targeting CAF-epithelial spatial interaction (the “high risk-epithelium” architectural unit), and at tissue level, targeting stromal homogeneity and dominance by Enlarged and Stellate myCAFs.

Scaling to patient resolution, we recapitulate these parallel axes and co-evolutionary processes by defining four unique and internally validated archetypes of PDAC, each characterised by a common microenvironment. These demonstrate prognostic performance exceeding conventional histopathological assessment, and predict response to adjuvant chemotherapy. While prior studies have defined CAF subtypes or architectural units based on local microenvironmental features^41,42^, this study is the first, to the best of our knowledge, to apply spatially resolved techniques to define architectural subtypes of PDAC. Notably, archetypes can be defined using just two proteins (CK and FAP), enabling integration into routine histopathological assessment. This demonstrates that with the addition of a limited spatial dimension, considerable prognostic and predictive gains can be made without reliance on high-plex spatial transcriptomics or proteomics.

In summary, we have demonstrated that hierarchical profiling of CAF biology and spatial architecture deciphers the complex stromal landscape of PDAC, reveals co-evolutionary axes, and defines architectural subtypes with high clinical utility. Together, these advances establish an accessible framework for spatial biomarker discovery to stratify treatment and guide therapeutic development in this devastating disease. Future work should extend morphological profiling to other cell types, particularly tumour epithelium, where epithelial subtype and cell shape have previously been linked^43,44^, and evaluate architectural subtyping across other cancers to test generalisability. Finally, machine-learning models trained on H&E alone could infer cell morphology and architectural subtypes without spatial proteomics, accelerating the translation of these findings to the clinic.

## Limitations

Our limited marker panel restricted proteomic resolution of CAF states. A considerable fraction of CAFs appeared proteomically “cold”, likely reflecting undefined CAF subtypes rather than true absence of markers, which may explain the broad range of survival correlations seen in the coldCAF tritype. Accordingly, we emphasised morphology and the iCAF-myCAF spectrum over highly granular CAF subtyping at proteomic level. In addition, our choice of FAP as a pan-CAF marker introduced potential under-labelling, whereby some cells tagged as “Other” cell may be FAP low/negative CAFs, rather than non-CAF stromal cells. This is consistent with our paradoxical observation of high Other cell density in the Organised archetype, while transcriptomic immune signals were most enriched in the Cohesive archetype. We therefore limited the interpretation of this unspecified cell population, and instead prioritised epithelial and CAF analyses.

## Methods

### Patients and tumour specimens

Our cohort included 227 patients who underwent curative surgery for primary PDAC from 2004 to 2013 inclusive. Cases were contributed by the Australian Pancreatic Cancer Genome Initiative (APGI – www.pancreaticcancer.net.au) (n = 201) and the West of Scotland Hepato-Pancreatico-Biliary Unit, Glasgow Royal Infirmary, Glasgow, UK (n = 26). No patients received neoadjuvant radiotherapy or chemotherapy. Pathological variables were harmonised to the AJCC 8^th^ edition. Recurrence patterns were defined by anatomical site at the first recurrence. Follow-up was maintained for a minimum of 24 months postoperatively to support accurate survival and recurrence analyses. Following review and annotation of whole-slide images, cases were spotted onto TMAs with a median of three cores per case to ensure multi-regional representation using methods described previously^45^.

### Ethical statement

The use of patient data and samples was approved by the UK NHS Health Research Authority Research Ethics Committee (reference 23/PR/0785) and by NHS Greater Glasgow & Clyde Research & Innovation (reference GN22ON553). Ethical approvals for APGI cases were obtained as previously described^45^.

### Immunofluorescence assays

The low-plex mIF assay was applied to all nine TMA slides (n = 227), whereas the high-plex assay was applied to a single TMA slide comprising UK cases only (n = 26). The high-plex slide was cut in serial section to ensure minimal spatial separation from the corresponding low-plex section. Both panels included core CAF characterisation markers (panCK, FAP, SMA, CTGF, S100A4, IL6, LIF, PDGFR) while the high-plex panel included additional markers for epithelial subtyping (FOXA1, GATA6, HNF4A, KRT17, P40, S100A2). Low-plex staining was performed on the Ventana Discovery Ultra (Roche Tissue Diagnostics) using Opal fluorophores, imaged on the PhenoImager HT, and spectrally unmixed using InForm software (all Akoya Biosciences). High-plex staining and imaging were performed integratively with the PhenoCycler Fusion platform (Akoya Biosciences). Comprehensive mIF protocols are provided in the Supplementary Methods. Following imaging, slides were washed and stained with haematoxylin and eosin (H&E) to enable pathology review and quality control.

### Image segmentation and quality-control

Images were imported into Visiopharm (Visiopharm A/S, version 2024.07.2.17212) for downstream spatial analysis using a custom segmentation pipeline. Nuclei were initially segmented using a deep neural network, followed by cytoplasmic segmentation based on thresholding of key placeholder markers to assign bulk cell types (CAF – FAP^POS^ CK^NEG^, Epithelial cell – FAP^NEG^ CK^POS^, Other cell – FAP^NEG^ CK^NEG^). A Laplacian transformation with a kernel size of 17 × 17 pixels was applied to FAP masks to delineate CAF boundaries.

The resulting dataset comprised 814 segmented TMA cores, each of which underwent manual quality control with reference to the corresponding H&E-stained core. Marker thresholds were assessed and, where necessary, adjusted to ensure concordance between immunofluorescence-based classification and H&E-based histomorphological ground truth. Regions of necrosis were manually excluded. Regions containing FAP^POS^ cells initially classified as CAFs but lacking characteristic CAF morphology were manually re-annotated as Other cell. Conversely, regions displaying clear epithelial morphology with aberrantly weak CK staining were manually re-annotated as Epithelial cell where necessary. Owing to the segmentation sequence (Epithelial cells classified prior to CAFs), misclassification of FAP^POS^ epithelial cells as CAFs was not possible. Overall, manual re-annotation was required for 3.2% of 2.5 million segmented cells.

In parallel with this process, entire cores were excluded if they met predefined exclusion criteria. For the low-plex assay, excluded cores comprised those with minimal remaining tissue (core largely or completely lost through sequential sectioning; n = 21), non-tumour tissue only (core composed entirely of muscle, fat, or necrotic tissue; n = 17), unsuitable annotations (secondary neuroendocrine tumour or normal pancreas control; n = 15), missing clinical data (n = 11), and technical artefacts (air bubbles or tissue folding/separation; n = 6). For the high-plex assay, excluded cores comprised those with technical artefacts (n = 7), minimal remaining tissue (n = 4), non-tumour tissue only (n = 4), and unsuitable annotations (n = 1). Following these exclusions, 643 low-plex and 85 high-plex cores were retained for downstream analysis.

### CAF feature normalisation and clustering

CAF phenotyping markers (SMA, CTGF, S100A4, IL6, LIF, and PDGFR) and geometric features (Area, PA ratio, Smoothness, Elongation, and Roundness) were initially Z-normalised within cohorts (APGI or UK) and subsequently censored at the 99^th^ percentile to mitigate the influence of extreme outliers. CAFs for which more than half of features required censoring were excluded (n = 155, <0.1%), as censoring of this extent collapses feature variance and would heavily bias downstream clustering. Leiden clustering was performed using the leiden_find_partition function from leidenbase (version 0.1.31) with iterative resolution_parameter and edge_weights as described (Fig. S2). Following clustering, morphotypes were defined by applying consensus clustering to the geometric feature centroid matrix of CAF subtypes using the ConsensusClusterPlus package (version 1.68.0) with parameters maxK = 4, reps = 1000, clusterAlg = “km”, and distance = “euclidean” (Fig. S3a).

Tritypes were defined as follows. For each CAF, a myCAF score (sum of SMA, CTGF, and S100A4 intensities) and iCAF score (sum of IL6, LIF, and PDGFR intensities) were computed, along with the cold pseudo-feature (Fig. S2h). These three scores were then aggregated by mean within each CAF subtype. Principal component analysis of these subtype-level means was then used to assign each of the fourteen CAF subtypes to a tritype (Fig. S3b).

Correlations between survival and raw CAF features were determined using mean feature intensity across all CAFs per patient, whereas those between survival and CAF subtype enrichment were determined by the proportion of each subtype among all CAFs per patient. The latter approach was also used to compute neighbourhood hazard ratios (Fig. 4a).

Intra-tumoural CAF heterogeneity was computed as follows. CAF subtype counts were first tabulated per TMA core, CLR-transformed (i.e. normalised per core), and Z-normalised by column (i.e. per CAF subtype). Pairwise Euclidean distances between each patient’s constituent TMA cores were then computed in this space, and the mean of these distances was defined as the ITCH score for that patient (Fig. S4).

### TME architecture scores

Spatial entropy was computed as previously defined^29^. Cohesion score was quantified by applying the DBSCAN algorithm^46^ to epithelial cell centroid coordinates using the dbscan package (version 1.2.0), with a minimum of one point per cluster. Based on a mean epithelial cell radius of 5µm, an epsilon distance of 10µm was used to infer spatial contact.

For each patient *p*, the cohesion score *Coh_p_* was defined as:

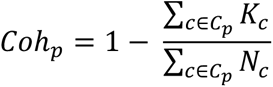

where 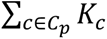 represents the total number of unique DBSCAN clusters across all tumour cores representing patient *p*, and 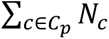 represents the total number of epithelial cells across those cores. Cohesion score therefore reflects the number of epithelial clusters normalised to the number of epithelial cells.

### Epithelial subtype scoring

Within the high-plex mIF cohort, intensities of canonical epithelial subtype markers were double z-normalised, and the raw subtype score was computed as the difference between the summed classical and summed squamous/basal-like marker signals. This score was then scaled to range from −1 to 1 to define the final subtype score.

### Other computational and statistical methods

All downstream computational analyses were performed in R (version 4.4.1). All plots were generated using the ggplot2 package (version 3.5.1), ggforce package (version 0.4.2), and with www.biorender.com. All nearest-neighbour computation was performed using the nn2 function from the RANN package (version 2.6.2). Delaunay triangulations for spatial gradient scoring were computed using the deldir package (version 2.0.4). Graph representations for spatial gradient scoring and Leiden clustering were computed using the igraph package (version 2.0.3). UMAP dimensionality reduction was performed using the uwot package (version 0.2.2). Survival analyses were performed using the survival package (version 3.6.4) and survminer package (version 0.4.9). Compositional data were CLR-transformed following zero imputation by multiplicative replacement using the clr function from the compositions package (version 2.0.8) and the cmultRepl function from the zCompositions package (version 1.5.0.4). For cross-validation and projection of new datasets, features were normalised using the mean and standard deviation of the training data, and samples were assigned to clusters by majority vote among the ten nearest neighbours in Euclidean space. Cross-validation folds were generated using the createFolds function from the caret package (version 6.0.94). Statistical significance was defined as p < 0.05. For visualisation, p values were denoted as follows: ns, not significant; •, p < 0.05; ••, p < 0.01; •••, p < 0.001.

### Genomic and bulk transcriptomic analyses

Existing genomic (n = 182) and bulk transcriptomic (n = 59) datasets comprised matched whole-genome or whole-exome sequencing and RNA sequencing (RNAseq) data from the APGI’s contribution to the International Cancer Genome Consortium (ICGC) Pancreatic Cancer project^23^. Data were downloaded from the ICGC Data Coordination Centre portal (release 28; 26 November 2019).

Following import into R, RNAseq data were converted into a DESeq2 object (package version 1.46.0). For patient DO33168, which had two RNAseq samples, sample SA410263 was retained for downstream analyses. Gene symbols were annotated using biomaRt (version 2.62.1) with reference to the GRCh37 genome build.

Transcriptomic subtypes were assigned using the ntp function from the CMScaller package (version 2.0.1) with parameters nPerm = 1000 and distance = “cosine”, using previously published reference gene signatures^23,31–33^. Gene-level expression values were used, retaining the gene with the highest mean log2 counts per million (CPM) where multiple Ensembl identifiers mapped to the same gene symbol. Count data were log2-transformed and quantile-normalised using normalizeBetweenArrays from the limma package (version 3.62.2) with parameters log2(counts + 0.25) and method = “quantile”, and subsequently Z-scaled prior to input into the ntp function.

For differential expression and GSEA, genes with insufficient expression, as determined by filterByExpr from edgeR (version 4.4.2), or lacking gene symbols were removed. DE analysis was performed using DESeq2 with a design matrix specified as ∼0 + Archetype, and differentially expressed genes were defined using an adjusted p-value threshold of 0.05. Volcano plots were generated using EnhancedVolcano (package version 1.24.0). Ranked log-fold changes were evaluated for pathway enrichment using the MSigDB Hallmark gene sets using the msigdbr package (version 25.1.1), and curated gene sets provided by the IOBR package (version 0.99.0)^30^, using the GSEA function from clusterProfiler (version 4.14.6). Gene sets were considered significantly enriched or depleted at an adjusted p-value threshold of 0.05.

For genomics analyses, where multiple samples were available from the same patient, a single sample was retained using the following prioritisation: primary tumour sample, metastatic sample, and then the sample with the highest tumour cellularity. Data were converted into a MAF object using the icgcSimpleMutationToMAF function from the maftools package (version 2.22.0). Duplicate variants were resolved by retaining the non-silent annotation where applicable. A total of 182 samples with matching mIF annotations were retained for downstream analyses. Oncoplots were generated using the oncoplot function in maftools. Copy number alterations were categorised as gains (copy number > 4) or losses (copy number < 2), excluding sex chromosomes, and structural variants were included where available. Driver genes were annotated as mutated if a sample harboured at least one non-silent mutation in the corresponding gene. Key PDAC driver genes were defined according to previously published criteria^47^.

## Author contributions

A.S.B.: conceptualisation, data curation, investigation, software, visualization, writing (original draft). S.M.: data curation, investigation, methodology. L.S.S.: formal analysis, investigation, methodology, software. L.O.J.: data curation, investigation, methodology, project administration, resources. R.L.B.: formal analysis, resources. F.B.: methodology. K.R.: methodology. I.R.P.: resources. D.T.: conceptualization. D.J.M.: conceptualization. C.W.: methodology. F.D.: validation. L.F.: methodology. A.C.Q.: methodology, software. K.Y.: methodology, software. J.L.Q.: conceptualization, funding acquisition, supervision. F.E.M.F.: conceptualization, funding acquisition, resources, supervision, writing (review & editing). S.B.D.: conceptualization, funding acquisition, supervision, writing (review & editing). D.K.C.: conceptualization, funding acquisition, resources, supervision, writing (review & editing).

## Acknowledgements

Australian Pancreatic Cancer Genome Initiative (APGI) for establishment and curation of tissue microarrays and associated clinical data. Glasgow Tissue Research Facility (GTRF) for support with slide preparation. NHS Greater Glasgow & Clyde Biorepository for storage, governance, and release of tissue material. Beatson Cancer Charity for support with the purchase of Visiopharm licenses. Catherine Ficken (Le Quesne lab) for technical support with laboratory work and slide scanning. Cancer Research UK Scotland Institute for support with technology infrastructure (via the Deep Phenotyping Advanced Technology Core Facility, RRID SCR_027366) and for access to high-performance computing resources.

## Data and code availability

Genomic and transcriptomic analyses use existing publicly available data, accessible at docs.icgc-argo.org/docs/data-access/icgc-25k-data. All original code has been deposited at <u>github.com/leonorss</u> and will be made publicly available as of the date of publication. Immunofluorescence images are subject to NHS Greater Glasgow & Clyde Biorepository ethical policies and a formal application and MTA are required for access. Further details, and any additional information required to reanalyse the data reported in this paper, are available from the lead contact upon request.

## Materials availability

This study did not generate new unique reagents.

---

**Fig. S1.**
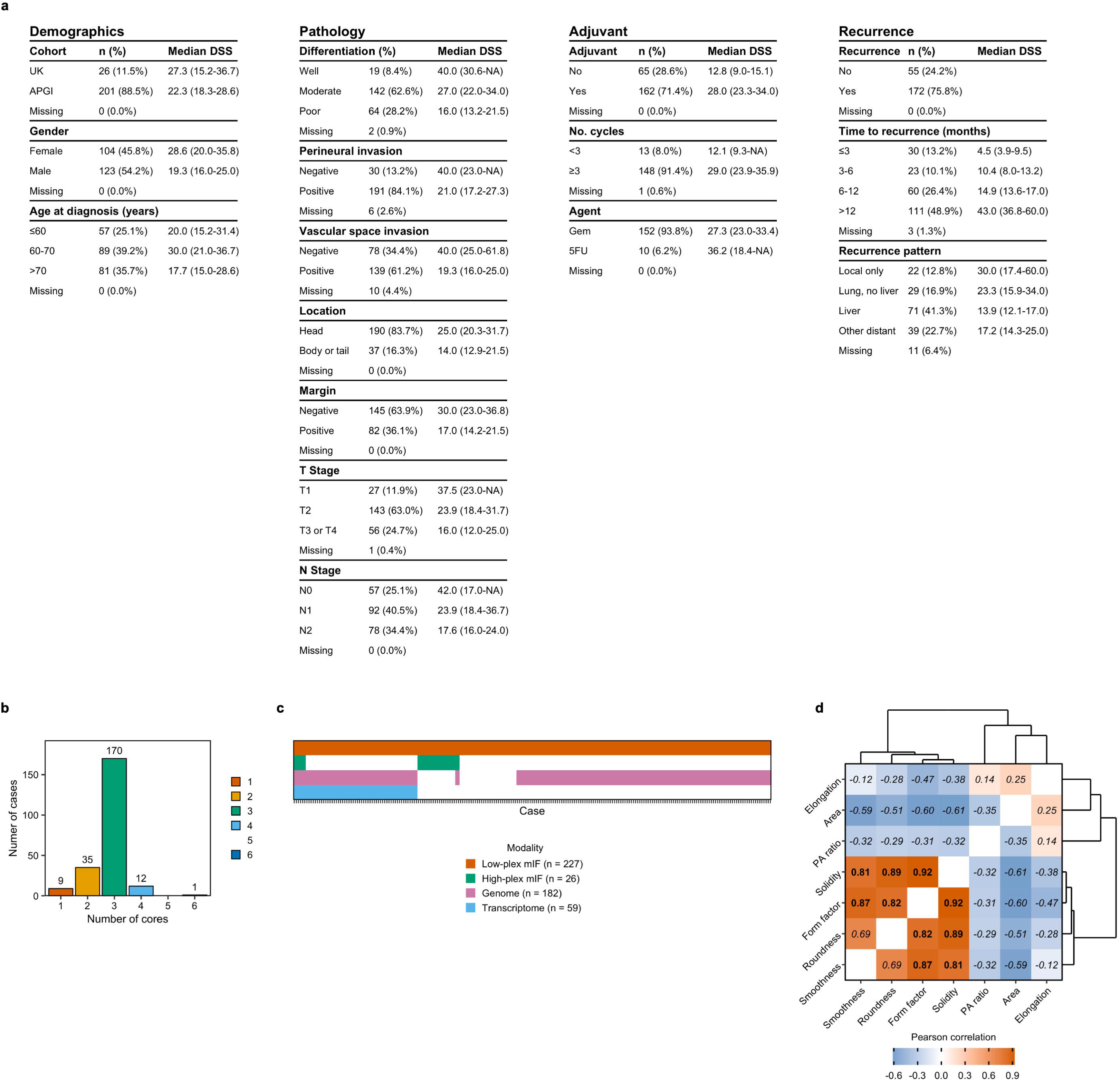
Summary of cohort and geometric features. **a,** Clinicopathological variables by count (%) and disease-specific survival (DSS) as median months (95% CI). **b,** Number of TMA cores per case. **c,** Overlap of modalities per case. **d,** Geometric features were filtered to minimise inter-feature correlation. Feature pairs with absolute Pearson correlation ≥ 0.7 were identified (bold) and one member of each pair was removed, yielding a final non-redundant set (italic).

**Fig. S2.**
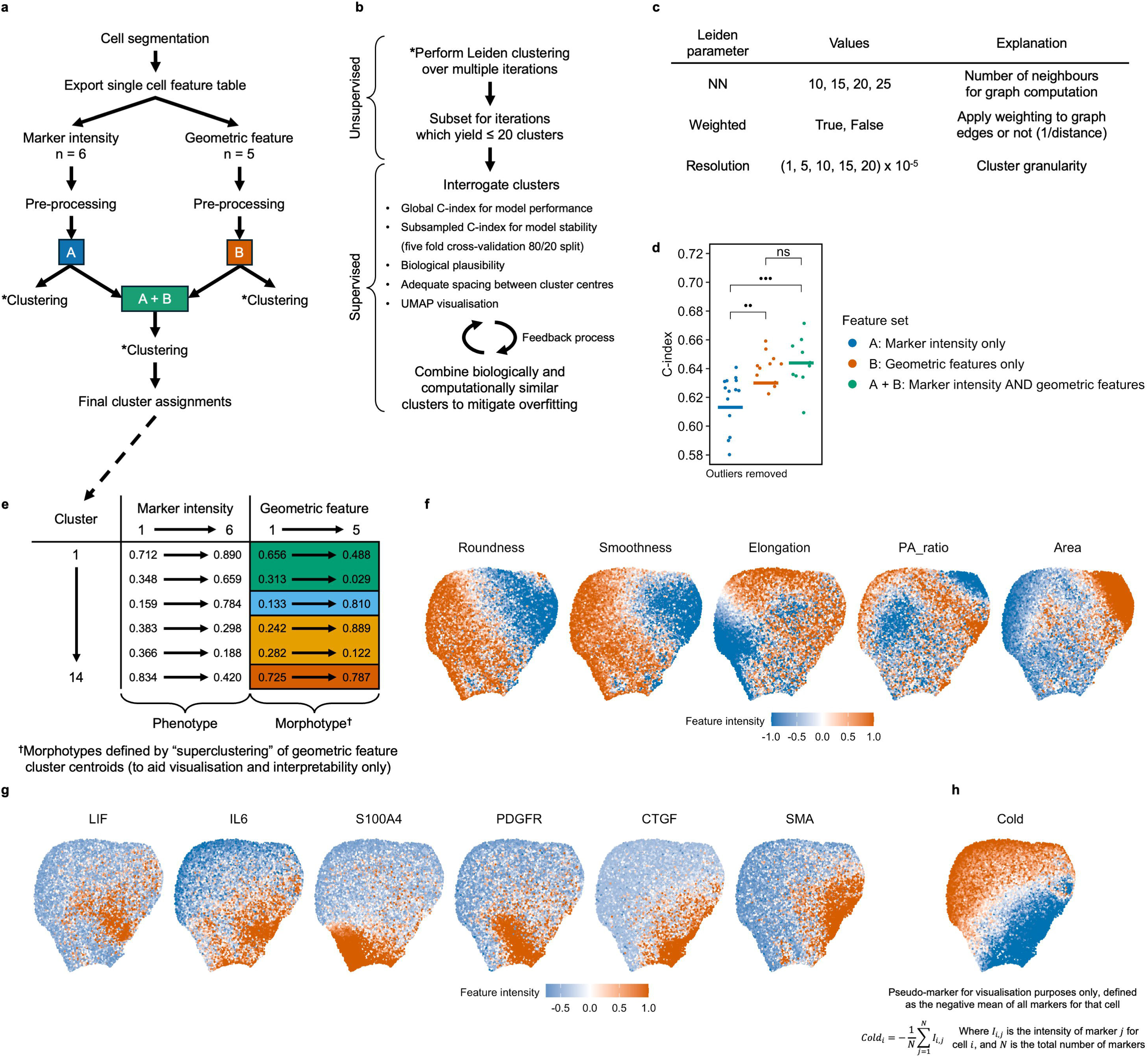
Defining CAF subtypes. **a,** Pre-processing workflow and overview of feature sets. **b,** Leiden clustering workflow demonstrating supervised and unsupervised steps. **c,** Clustering was iteratively performed across multiple nearest neighbour, weighting, and resolution parameters. **d,** Integrating geometric and proteomic feature sets yielded greater prognostic utility than each set in isolation. Crossbar = mean. Comparison = Wilcoxon test. **e,** Example cluster centroid table and morphotype assignments. **f-g,** UMAPs false coloured by feature intensity. **h,** Computation of the “cold” pseudo-feature.

**Fig. S3.**
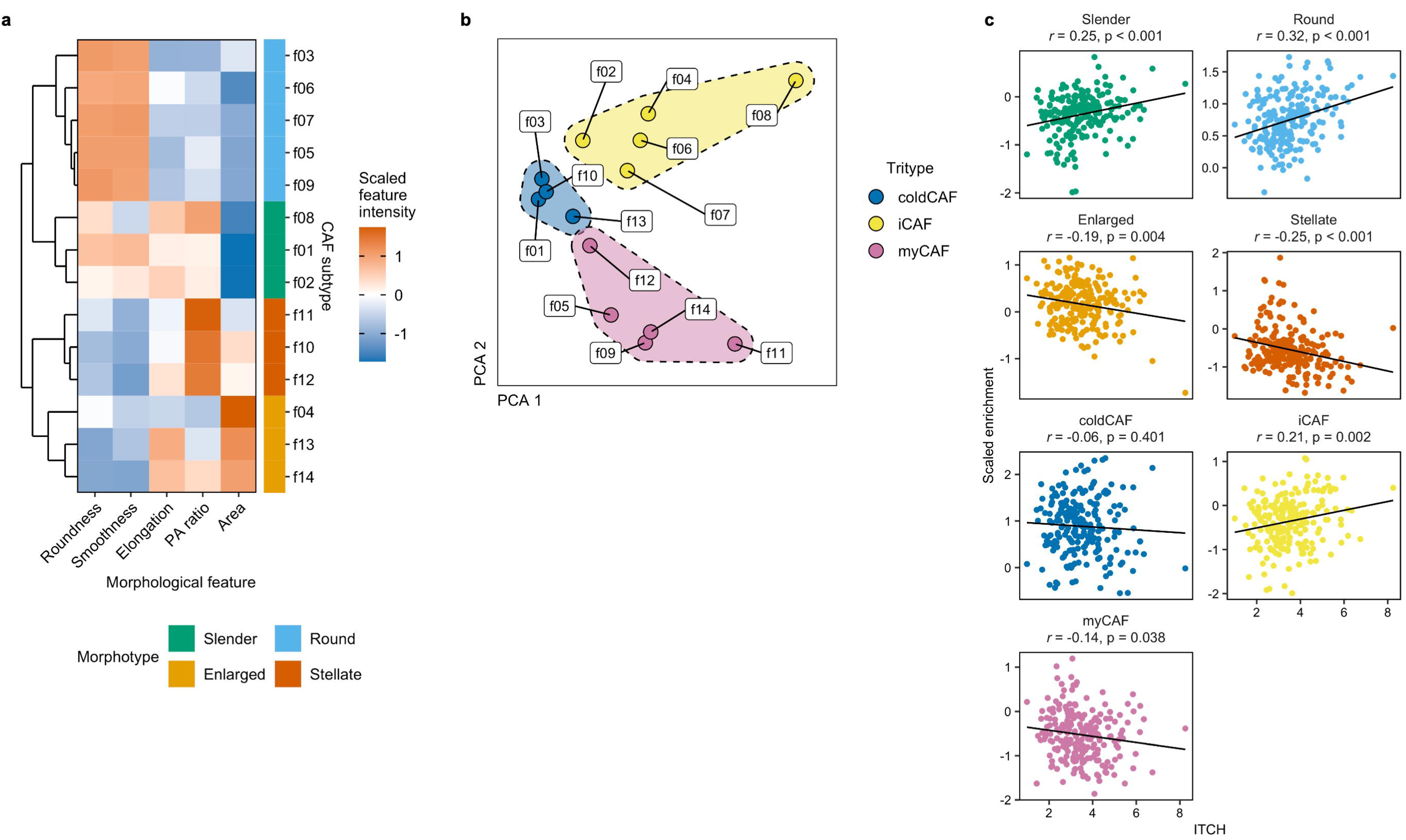
Defining morphotypes and tritypes. **a,** Morphotypes were defined by clustering mean morphological feature intensity (cluster centroids) for each CAF subtype. **b,** Tritypes were defined by clustering coldCAF, iCAF, and myCAF scores for each CAF subtype. **c,** Pearson correlations between ITCH and CAF morphotype and tritype enrichment.

**Fig. S4.**
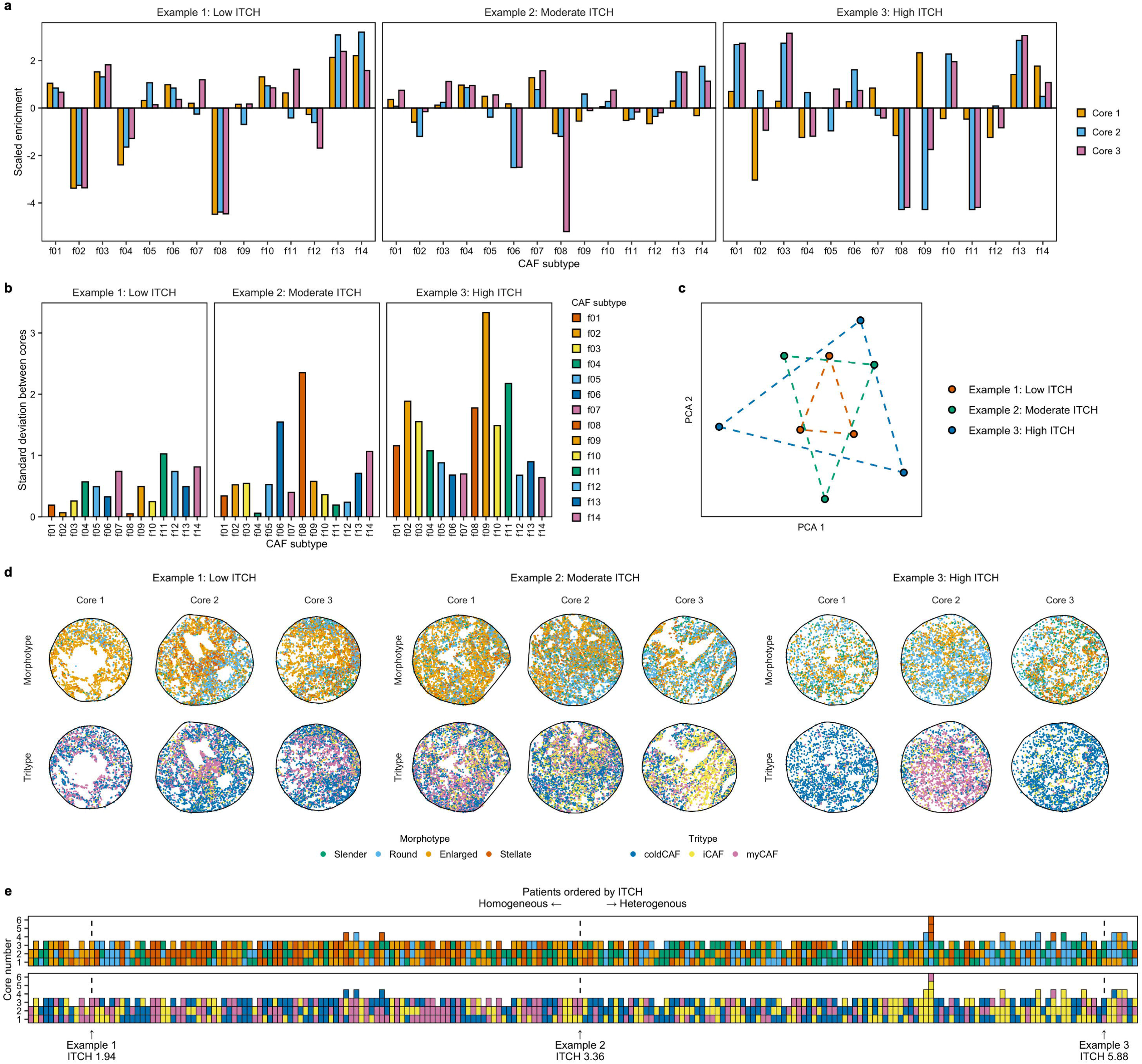
Quantifying intra-tumoural CAF heterogeneity (ITCH). **a,** CAF subtype enrichment in three example cases (low, moderate, and high ITCH). **b,** Standard deviation of CAF subtype enrichment between cores. **c,** Dimensionality-reduced CAF subtype enrichment (one point represents a single TMA core). **d,** Visualisation of each core. **e,** Each tile (representing a single TMA core) is coloured by dominant morphotype and tritype to highlight the differences in CAF subtype enrichment between homogeneous and heterogeneous tumours.

**Fig. S5.**
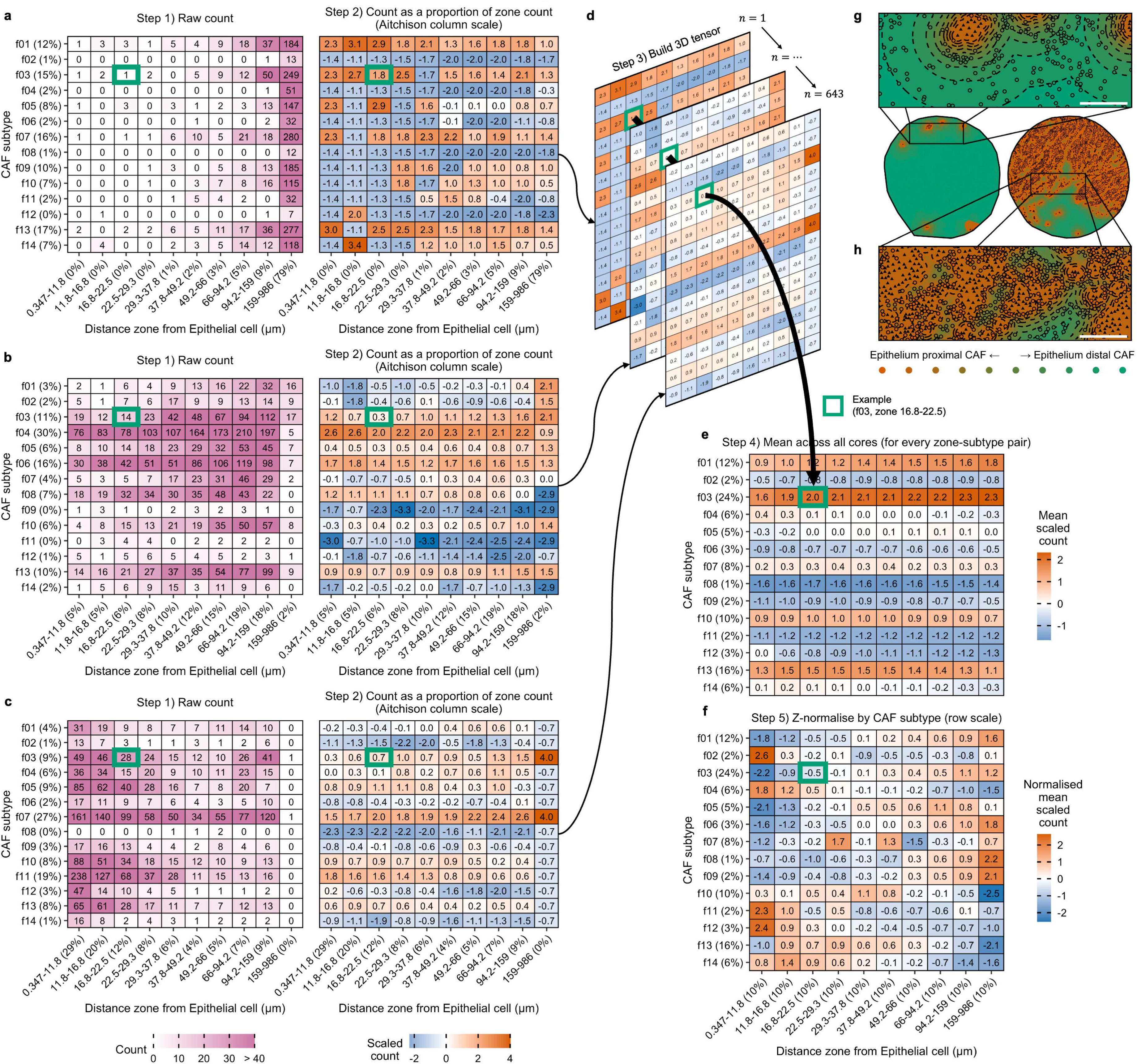
Normalising spatial correlations I (proximity analysis pipeline). **a-c,** Raw and normalised CAF subtype counts per distance zone in example low- (**a**), mid- (**b**), and high-(**c**) Epithelial cell density cores. **d,** Matrices were assembled in to a tensor. **e,** Means were computed by tensor rotation. **f,** Final normalisation. Step 2 corrects for anchor cell density, while step 5 corrects for CAF subtype prevalence. **g-h,** Cores from which tables **a** and **c** are derived, highlighting the potential for differences in zone counts to be a random effect of varying anchor cell density. Dashed lines = distance zone boundaries. Solid triangles = epithelial cells. Scale bar = 100μm.

**Fig. S6.**
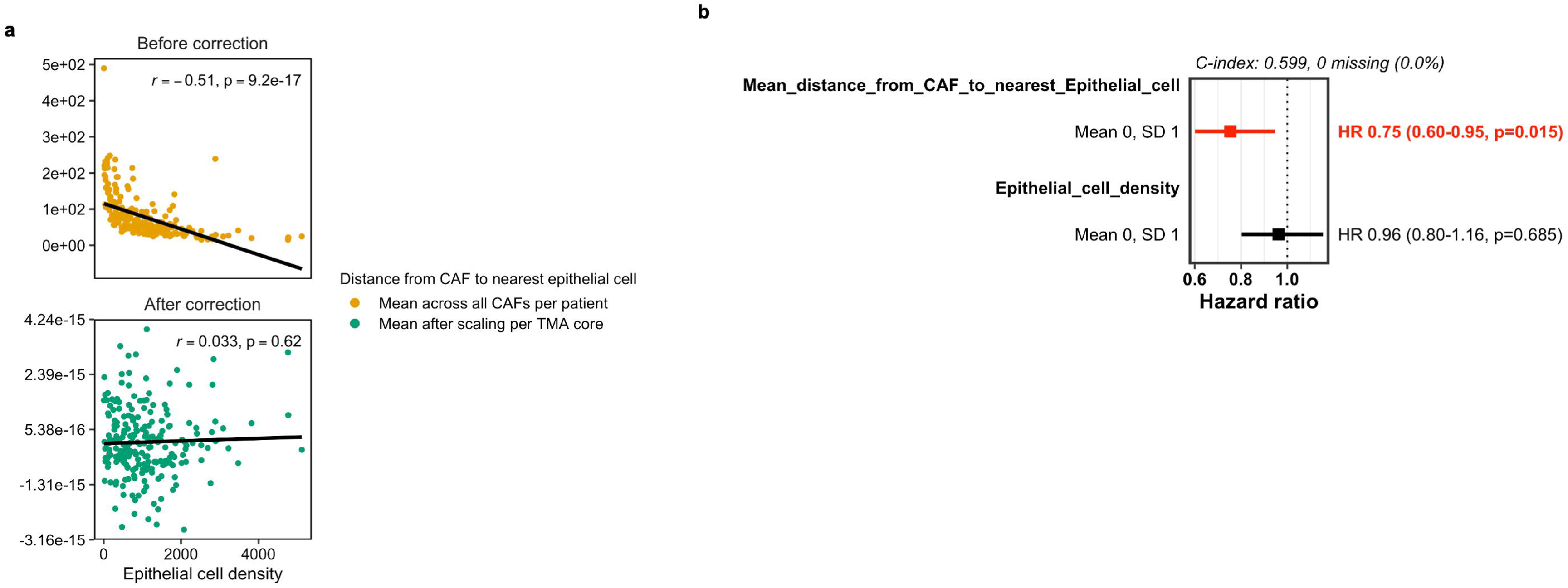
Normalising spatial correlations II and III. **a,** Method II breaks the relationship between anchor cell density and distance to anchor cell by normalising distances at core level. *r* = Pearson correlation. **b,** Survival correlations with distance to anchor cell (method III) remain significant after accounting for anchor cell density.

**Fig. S7.**
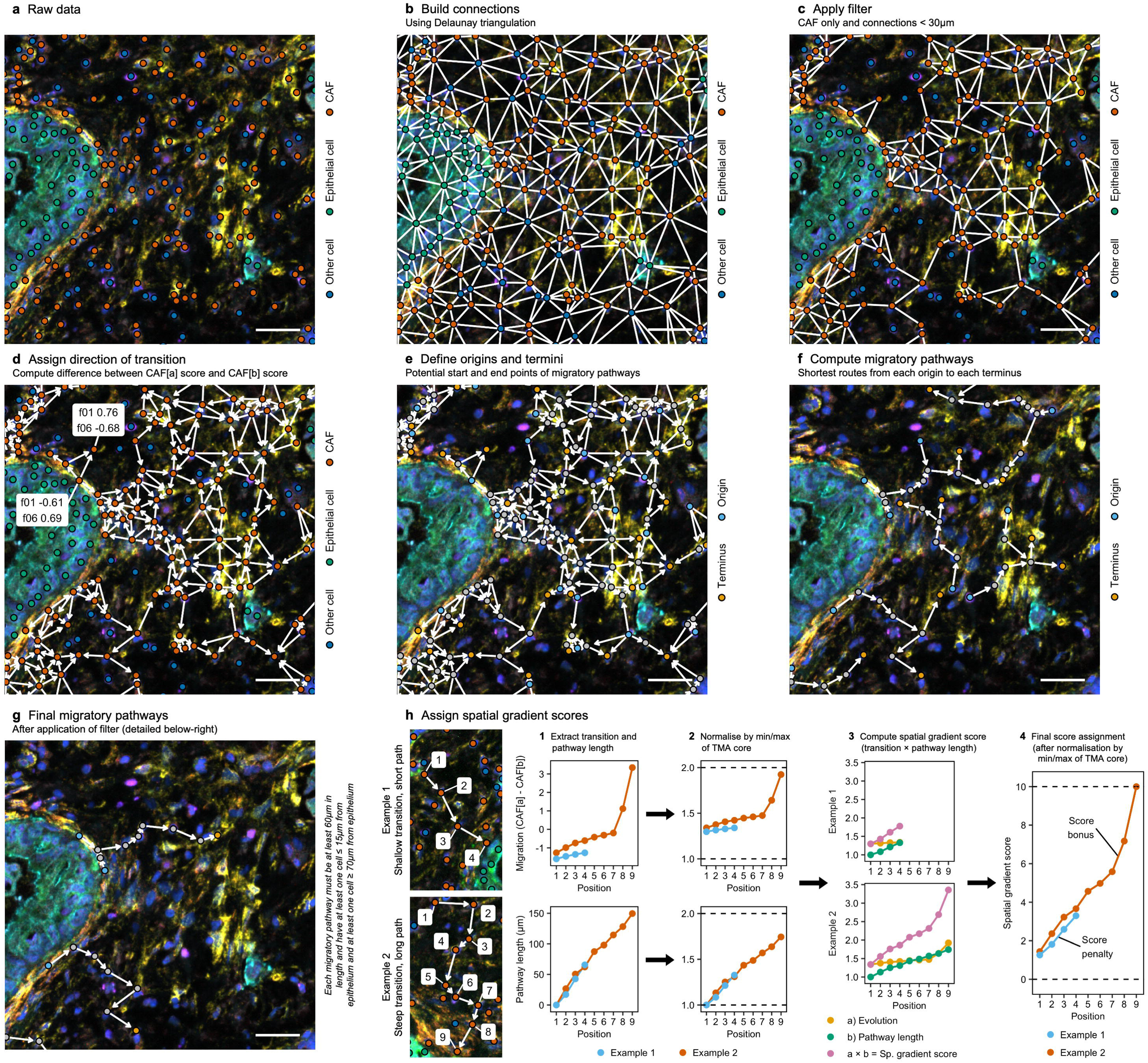
Spatial gradient score pipeline. **a,** Raw data. **b,** Unlike nearest neighbour approaches, Delaunay triangulation only connects cells with a direct “line of sight”. **c,** Only connections < 30µm were deemed to be spatially related. **d,** Directions of transition were assigned for each CAF subtype pair (f01 > f06 shown for example). **e,** Origin cells (all connections directing away) and termini cells (all connections directing towards) were defined. **f-g,** Shortest paths for each origin-terminus pair were computed using Dijkstra’s algorithm and subsequently filtered. **h,** Each cell was assigned a spatial gradient score, ensuring long paths with a steep transition gradient are rewarded, and short paths with a shallow transition gradient are penalised. Steps **d**-**g** were then repeated for each CAF subtype pair. The entire pipeline was then repeated separately for each core. Scale bar = 30µm.

**Fig. S8.**
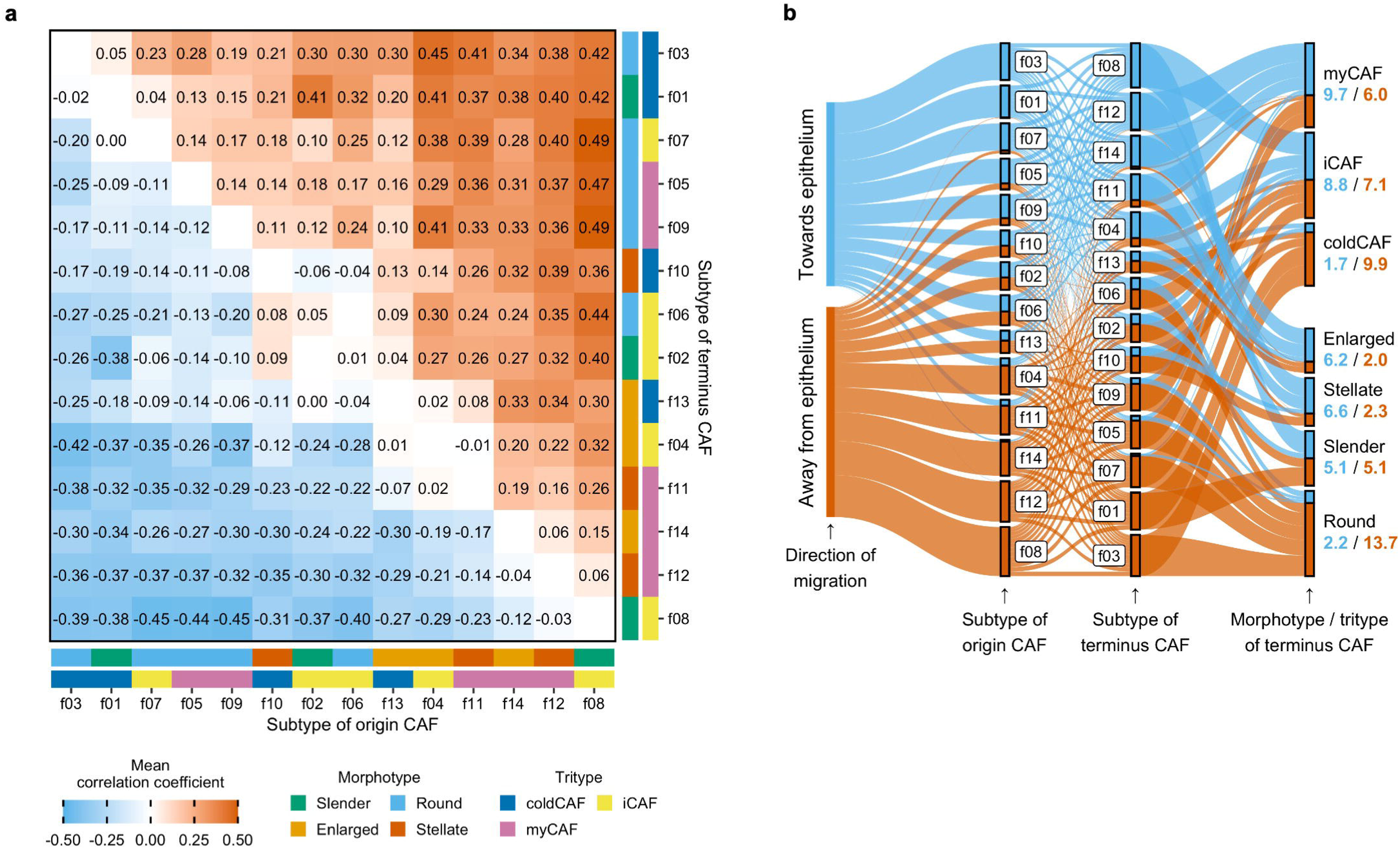
Spatial gradient score trajectories. **a-b,** Pearson correlation coefficients for each subtype transition by mean (heatmap, **a**) and summed mean (Sankey diagram, **b**). As spatial gradient score rises, distance to epithelium falls for blue pairs (inferred trajectories towards epithelium), and rises for orange pairs (inferred trajectories away from epithelium).

**Fig. S9.**
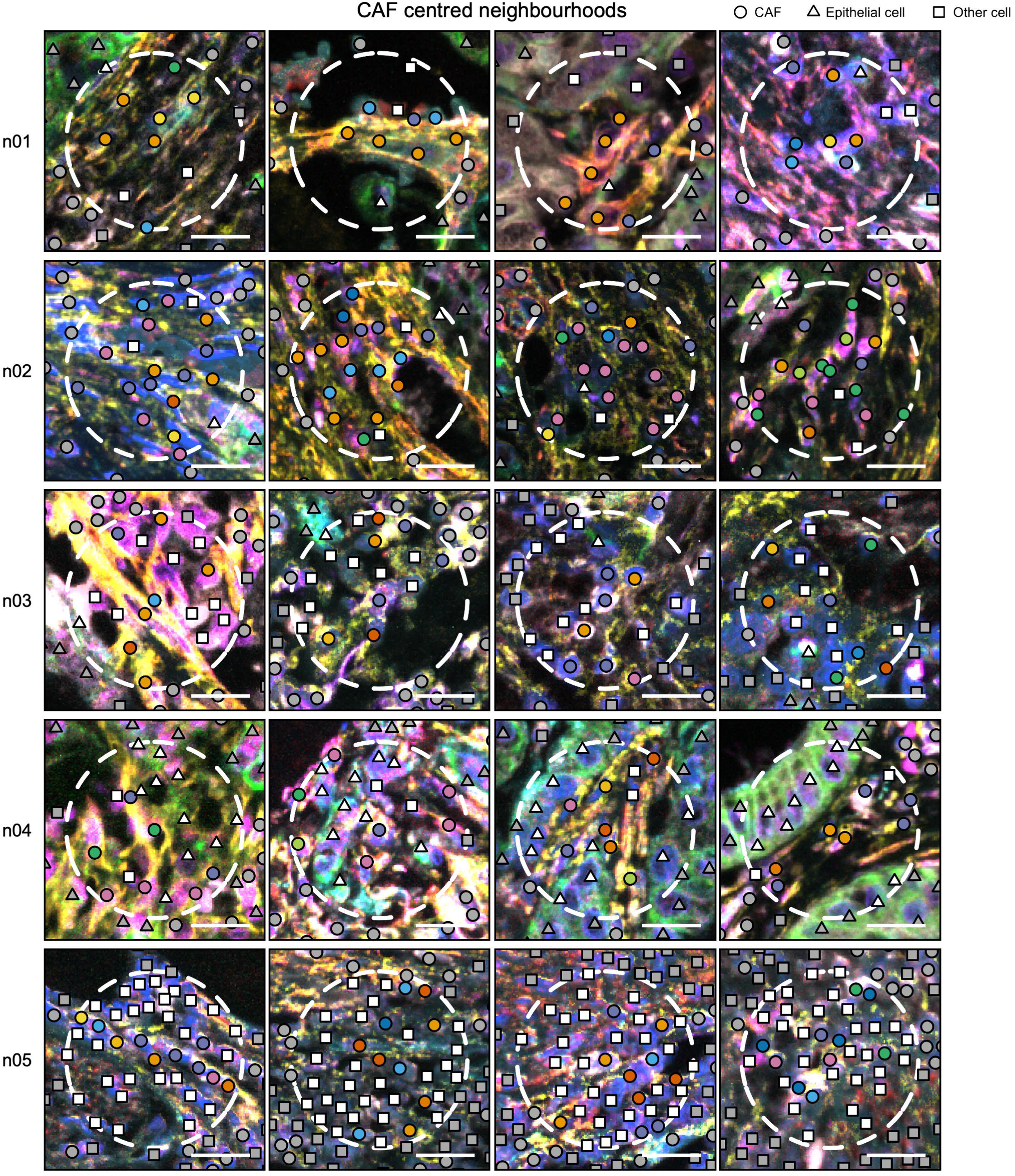
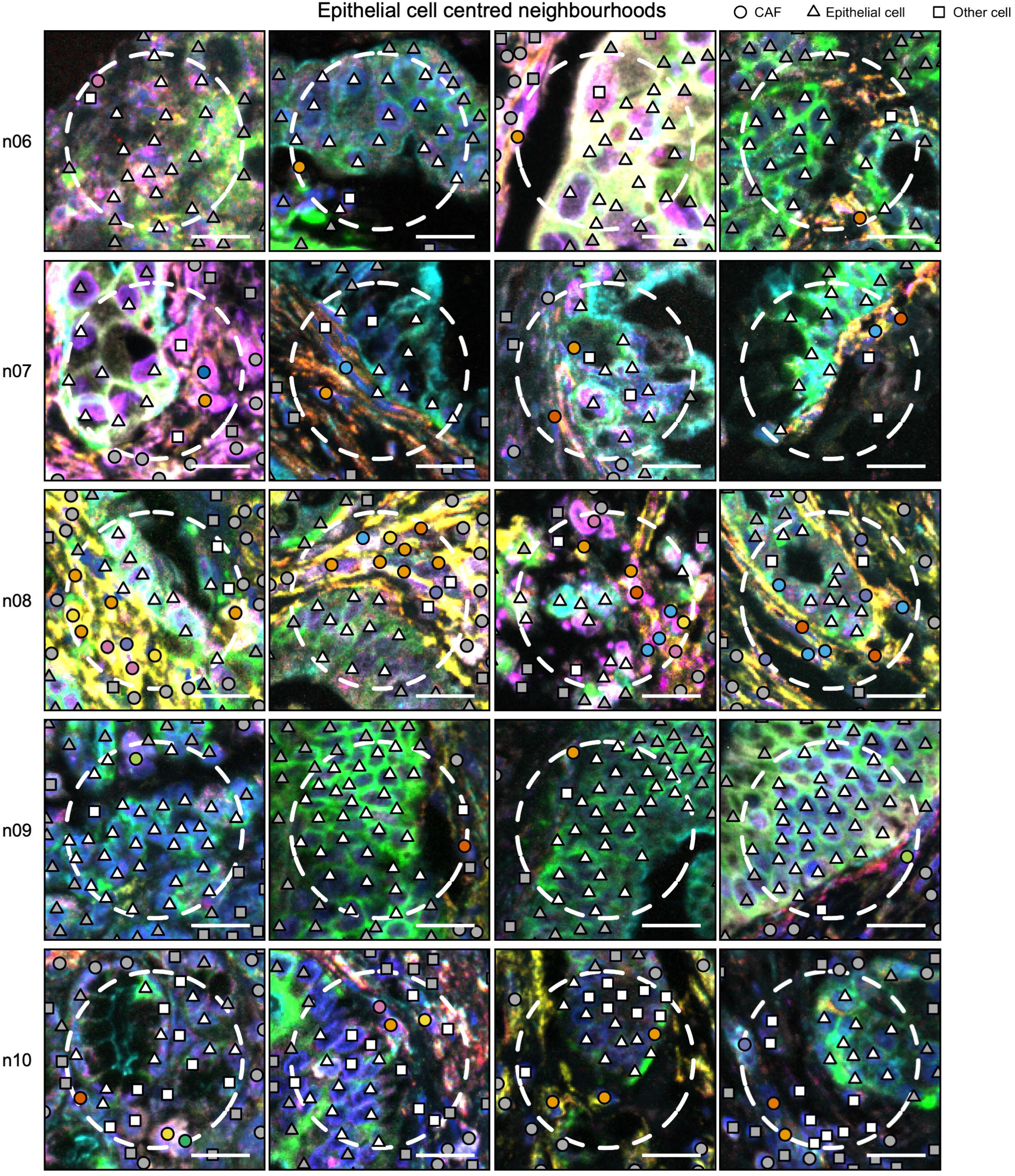
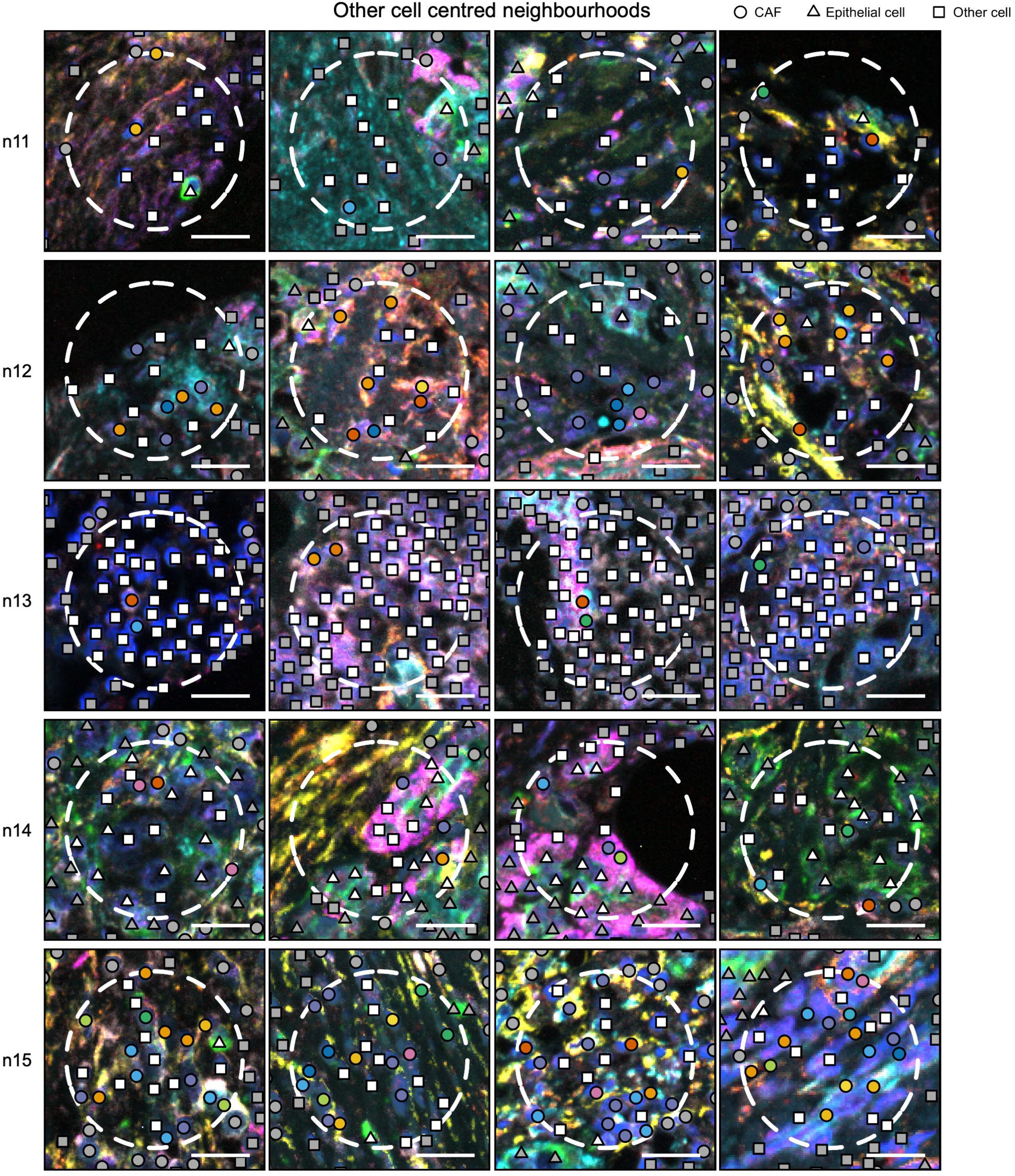
Representative neighbourhood examples. Neighbourhood radius = 30µm. Scale bar = 20µm.

**Fig. S10.**
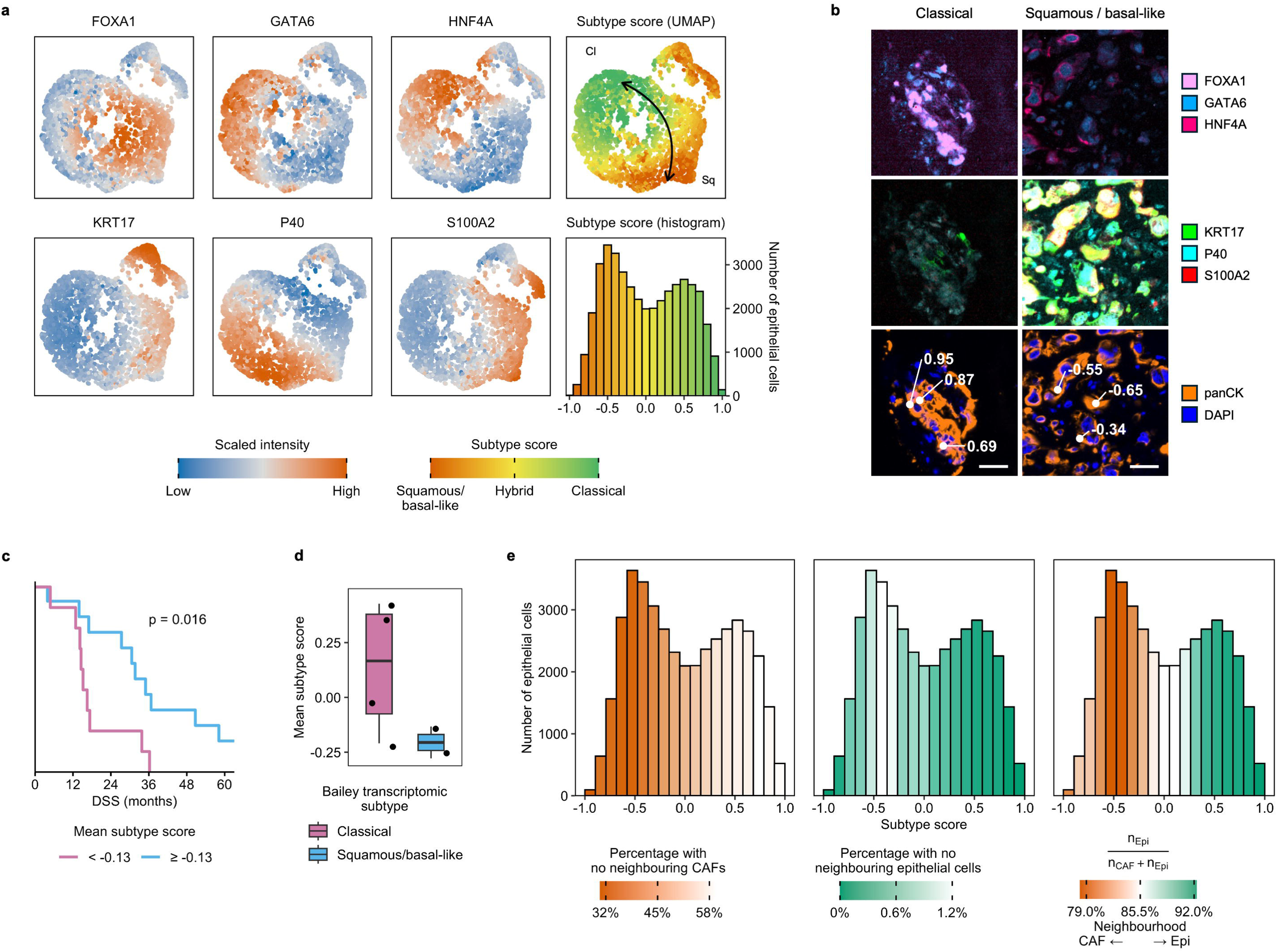
Epithelial subtype scoring. **a,** Epithelial cell UMAP (high-plex mIF cases only) coloured by classical (top row) and squamous/basal-like (bottom row) markers, and by subtype score with line of best fit. **b,** Example epithelial cells with annotated subtype score (bottom row). **c,** High mean subtype score correlated with survival (lower 40% squamous/basal-like, upper 60% classical). p = Log-rank test. **d,** Subtype score correlated with previously defined bulk transcriptomic epithelial subtype. **e,** Classical epithelial cells were more frequently found with no neighbouring CAFs, whereas squamous/basal-like epithelial cells were more frequently isolated from neighbouring epithelial cells.

**Fig. S11.**
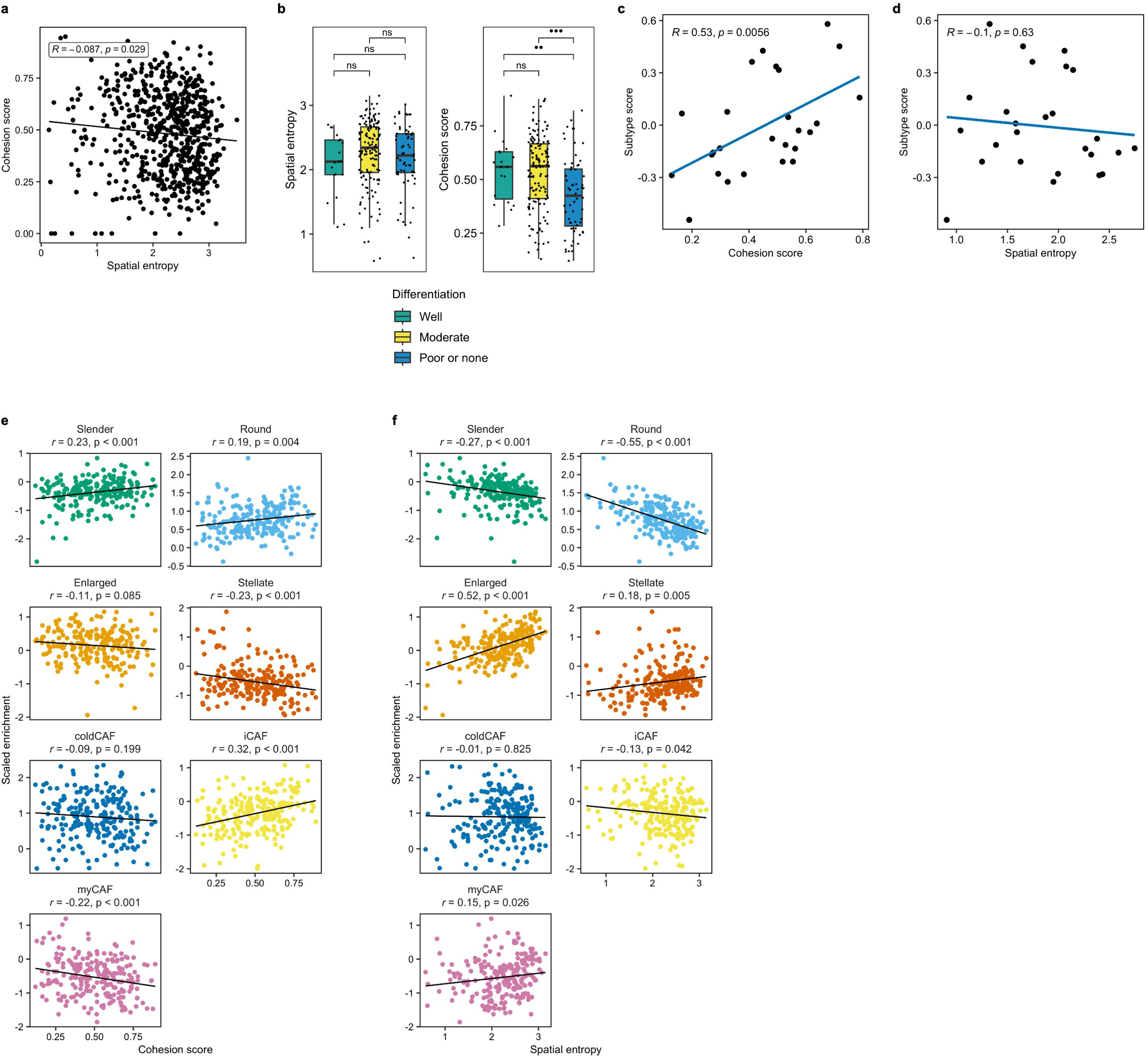
Quantifying global differences in TME architecture. **a,** Pearson correlation between cohesion score and spatial entropy. **b,** Relationships between tumour differentiation and spatial entropy / cohesion score. Comparison = Wilcoxon test. **c-d,** Pearson correlation between subtype score and cohesion score (**c**) and spatial entropy (**d**). **e-f,** Pearson correlations between CAF subtype enrichment and (**e**) cohesion score and (**f**) spatial entropy.

**Fig. S12.**
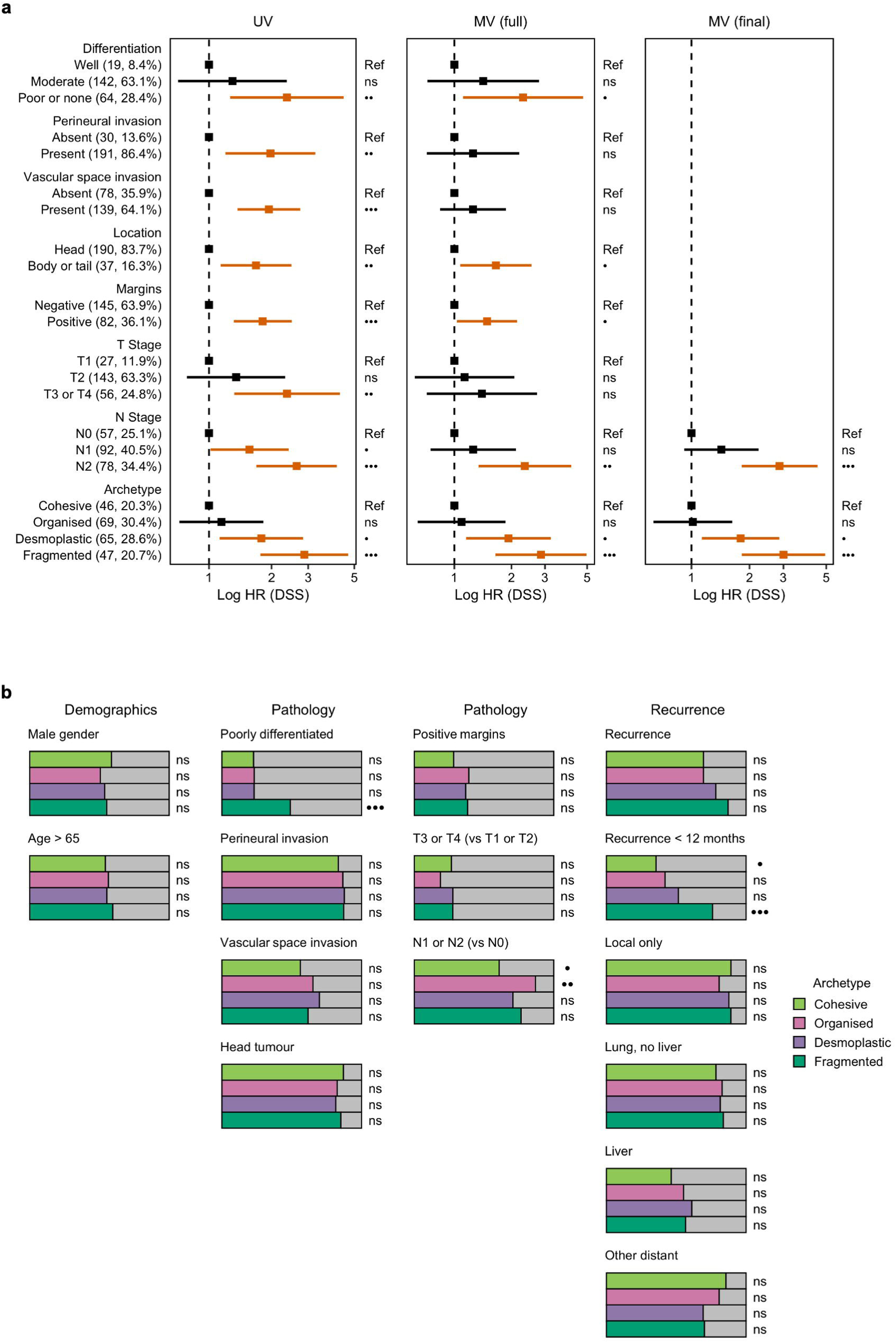
Relationships between archetypes and clinical features. **a,** Multivariate Cox regression model demonstrating the independent prognostic utility of archetypes (backwards elimination with significance threshold of 0.01). **b,** Relationships between archetypes and clinicopathological features. p = Fisher test.

**Fig. S13.**
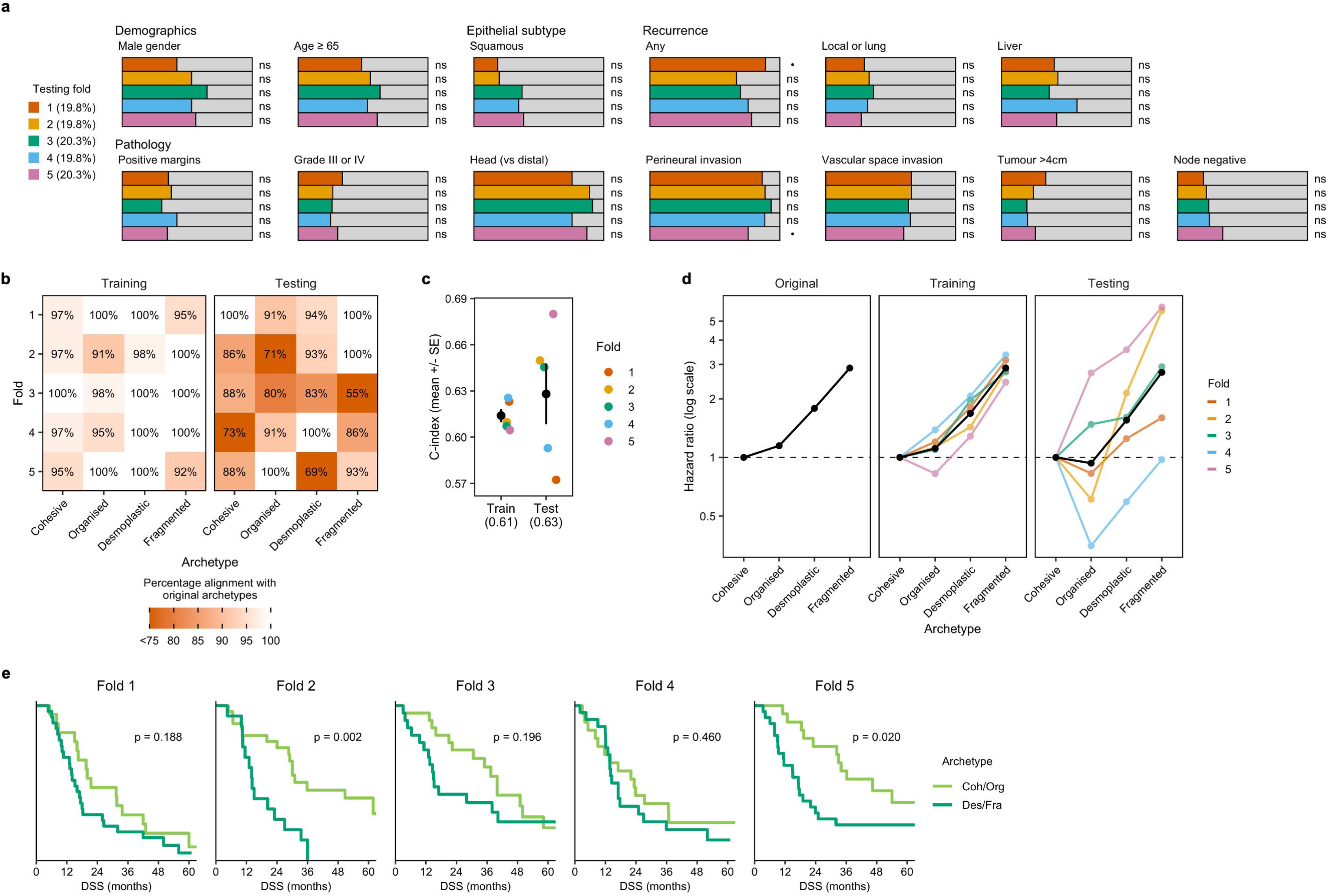
Internal validation of archetypes. **a,** Histopathological parameters are approximately equivalent across folds. p = Fisher test. **b,** Alignment of fold archetypes with original archetypes (percentage per fold). **c,** Mean train and test c-indices. **d,** Archetype univariate hazard ratios by fold, with mean per archetype (black line). **e,** Testing-only survival plots (archetypes reduced to two groups). p = Log-rank test.

**Fig. S14.**
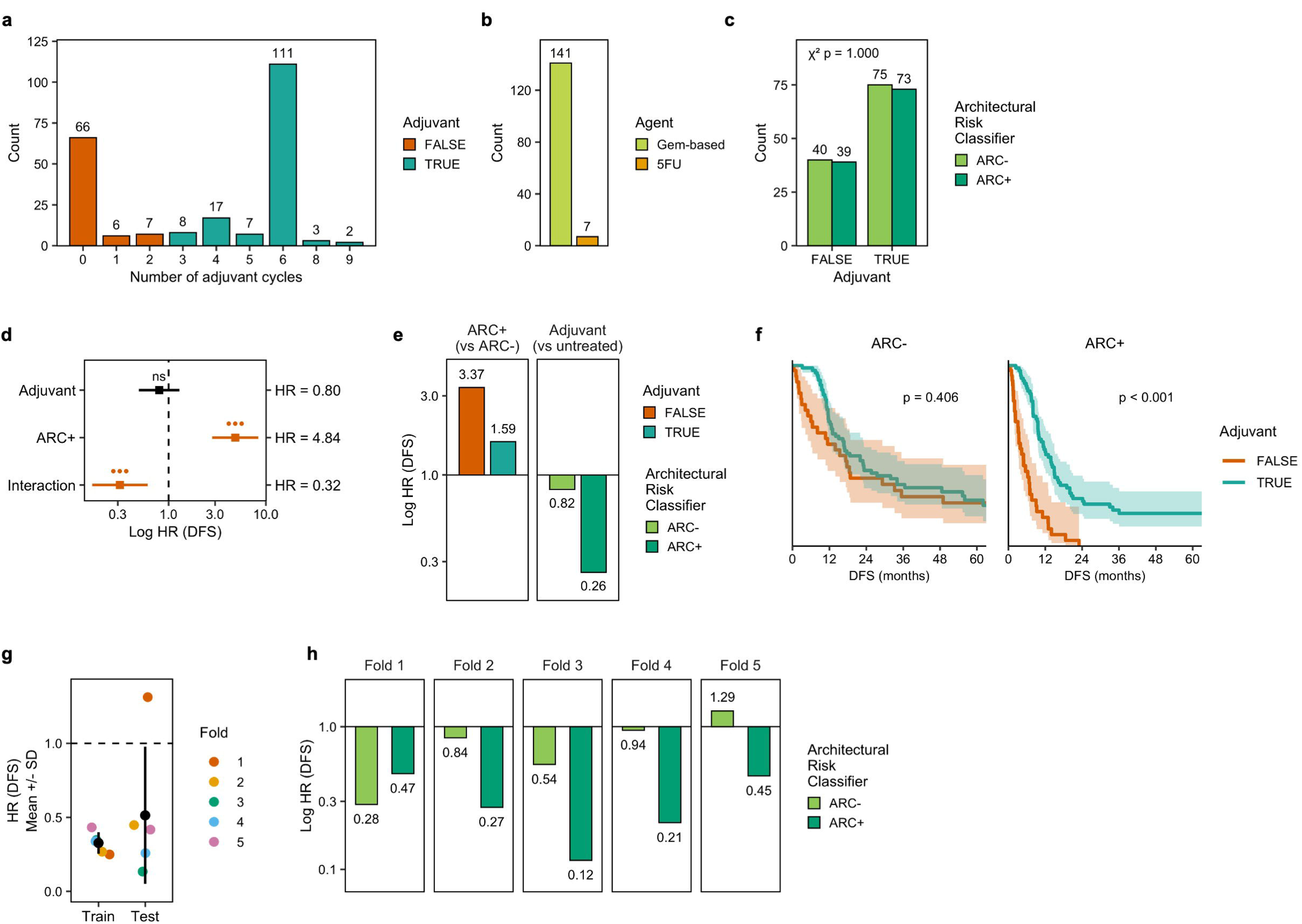
Predictive utility of architectural subtyping. **a-b,** Summary of adjuvant chemotherapy cycles and regimens. Only patients completing at least three cycles of treatment were considered to have received adjuvant chemotherapy. **c,** ARC status and adjuvant treatment are not correlated. **d,** Interaction between ARC status and adjuvant treatment. **e,** Hazard ratios of adjuvant treatment by ARC status, and ARC status by adjuvant treatment. The significant hazard reduction between ARC- patients receiving adjuvant and ARC+ patients receiving adjuvant highlights the predictive utility of the ARC classifier. **f,** Survival by ARC status, with and without adjuvant treatment. p = Log-rank test. **g,** Adjuvant × ARC interaction HRs between train and test folds demonstrate the stability and generalisability of the ARC classifier. **h,** Reduction in hazard by ARC status in patients who received adjuvant treatment (testing folds only).

**Fig. S15.**
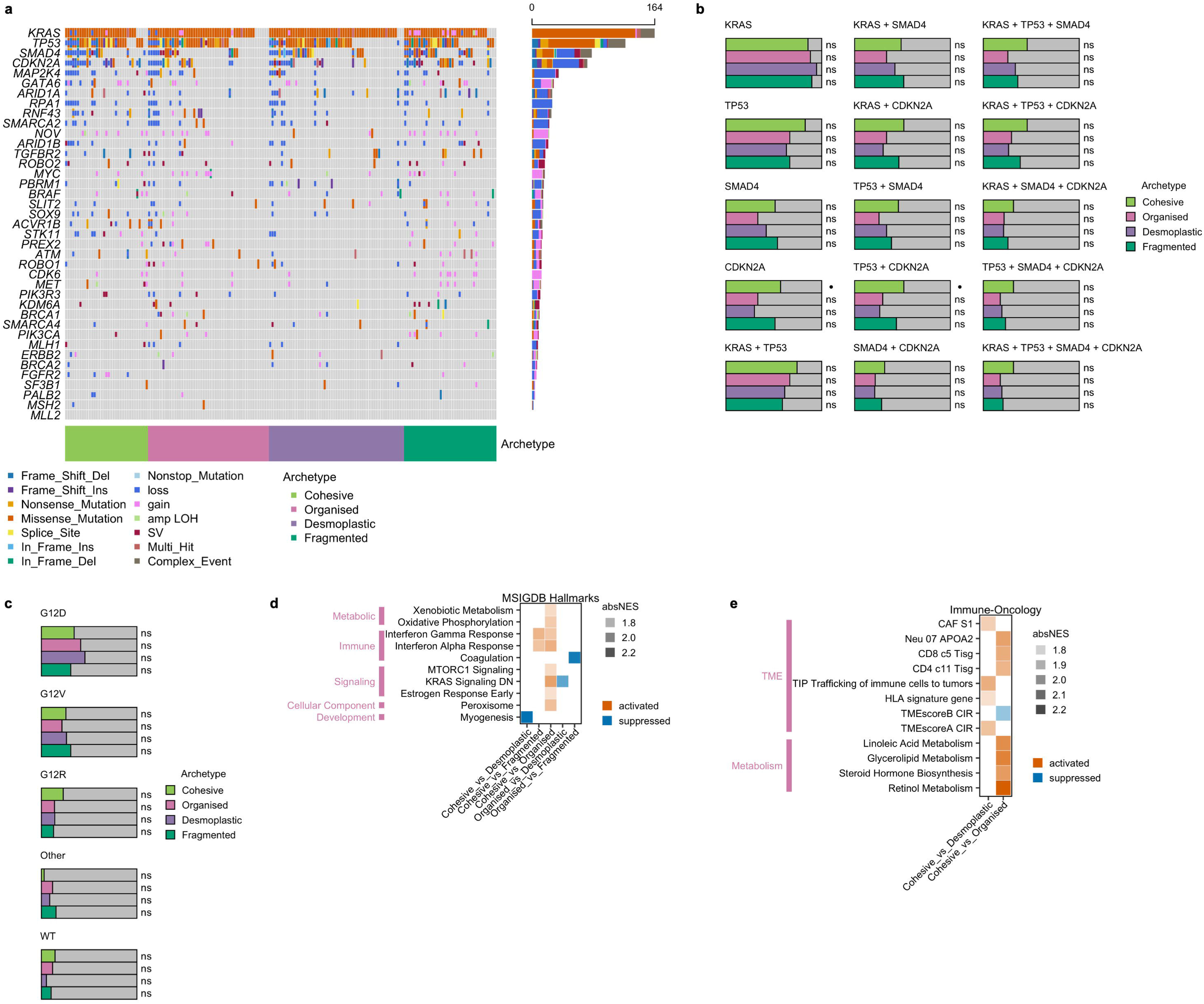
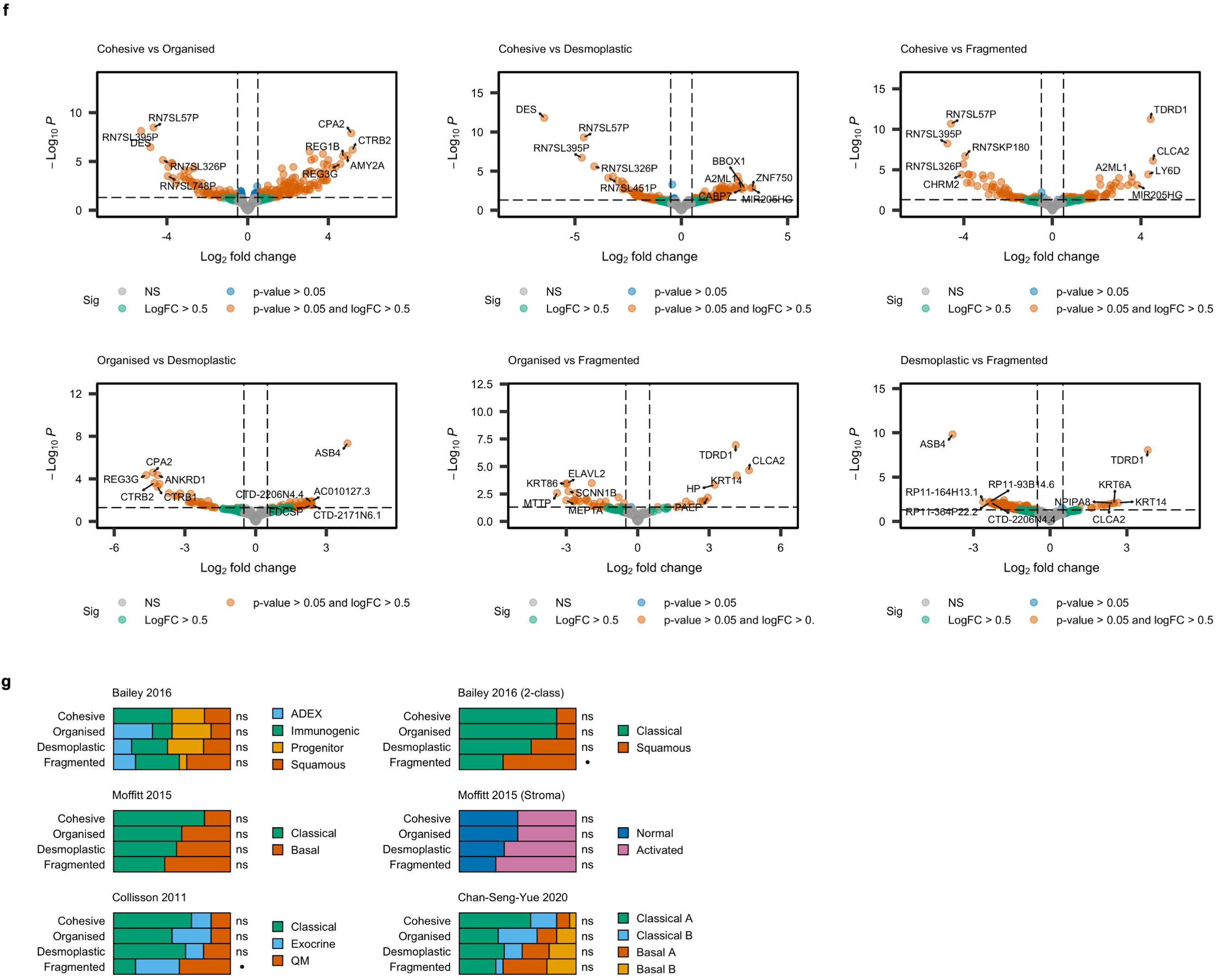
Architectural subtypes map to molecular programmes. **a,** Oncoplot demonstrating mutational burden across archetypes in key PDAC genes. **b,** Groupwise comparisons of mutational burden by archetype (top four genes only, mutated vs wild type). p = χ^2^ test. **c,** KRAS variant status by archetype. p = χ^2^ test. **d-e,** Gene set enrichment analysis with MSigDB hallmarks (**d**) and IOBR signatures (**e**). **f,** Volcano plots demonstrating differentially expressed genes across archetype pairs. **g,** Enrichment of canonical transcriptomic programmes across archetypes. p = Fisher test.

