## Supplementary Methods for "Morphospatial profiling of cancer-associated fibroblasts reveals architectural subtypes of pancreatic ductal adenocarcinoma"

*Low-plex immunofluorescence assay*

The low-plex mIF assay was applied to all nine TMA slides. All slides were stained in one batch and positive and negative controls were stained prior to the staining run to ensure the assay was performing as expected. All staining was fully automated on the Ventana Discovery Ultra (RUO Discovery Ultra, Roche Tissue Diagnostics, version 21.00.0019). Antibodies had been previously validated on known positive control tissue and on pancreatic carcinoma control tissue in both chromogenic DAB immunohistochemistry (IHC) and fluorescence individually. Sections were cut at 4µm on TOMO slides (Ref. TOM-1190) and were baked for 1 hour at 60 degrees prior to loading onto the Ventana Discovery Ultra. All other staining steps were performed on the instrument.

Sections were dewaxed and pretreated with Discovery CC1 (Roche Tissue Diagnostics, ref. 06414575001) for 32 minutes at 95°C. A stripping step was performed after each staining sequence using CC2 (Roche Tissue Diagnostics, ref. 05279798001) at 100°C.

The antibody-fluorophore sequences were applied in the following order: Anti-CTGF (ABCAM, ref. ab6992) was applied at 1/25 for 32 minutes at 37°C followed by Omnimap anti-rabbit HRP (Roche Tissue Diagnostics, ref. 05269679001) incubated for 24 minutes, detected by Opal 570 (Akoya Biosciences, ref. FP1488001KT) at 1/100 for 8 minutes. Anti-FSP1/S100A4 (Sigma-Aldrich, ref. 07-2274) was applied at 1/25 for 32 minutes at 37°C followed by Omnimap anti-rabbit HRP (Roche Tissue Diagnostics, ref. 05269679001) incubated for 12 minutes, detected by Opal 650 (Akoya Biosciences, ref. FP1496001KT) at

1/100 for 8 minutes. Anti-SMA (clone 1A4) (Roche Tissue Diagnostic, ref. 05268303001) was applied ready-to-use, incubated for 32 minutes at 37°C followed by Omnimap anti-mouse HRP (Roche Tissue Diagnostics, ref. 05269652001) for 12 minutes, detected by Opal 540 (Akoya Biosciences, ref. FP1494001KT) at 1/200 for 8 minutes. Anti-PDGF receptor alpha (clone D1E1E) (Cell Signalling Technology, ref. 3174) was applied at 1/1000 for 32 minutes at 37°C followed by Omnimap anti-rabbit HRP (Roche Tissue Diagnostics, ref. 05269679001) incubated for 12 minutes, detected by Opal 620 (Akoya Biosciences, ref. FP1495001KT) at 1/50 for 8 minutes. Anti-IL-6 (Clone 1.2-2B11-2G10), (ABCAM, ref. ab9324) was applied at 1/25 incubated for 32 minutes at 37°C followed by Omnimap anti-mouse HRP (Roche Tissue Diagnostics, ref. 05269652001) for 12 minutes, detected by Opal 690 at 1/100 for 8 minutes. Anti-Fibroblast activation protein, alpha (clone EPR20021), (ABCAM, ref. ab271976) was applied at 1/25 incubated for 90 minutes at room temperature, followed by Ultramap anti-rabbit HRP (Roche Tissue Diagnostics, ref. 05269717001) for 24 minutes, detected by Opal 520 (Akoya Biosciences, ref. FP1487001KT) at 1/100 for 8 minutes. Anti-LIF (Proteintech, ref. 26757-1-AP) was applied at 1/200 for 32 minutes at 37°C followed by Omnimap anti-rabbit HRP (Roche Tissue Diagnostics, ref. 05269679001) incubated for 12 minutes, detected by Opal 480 (Akoya Biosciences, ref. FP1500001KT) at 1/25 for 8 minutes. Anti-Cytokeratin (clone AE1/AE3) (Leica, ref. NCL-L-AE1/AE3-601-L-CE) was applied at 1/250 for 28 minutes at 37°C followed by Omnimap anti-mouse HRP (Roche Tissue Diagnostics, ref. 05269652001) for 12 minutes, detected by TSA-DIG (Akoya Biosciences, ref. FP1502001KT) at 1/100 for 12 minutes followed by Opal 780 (Akoya Biosciences, ref. FP1501001KT) at 1/10 for 1 hour.

QD DAPI (Roche Tissue Diagnostics, ref. 05268826001) was applied as a counterstain for 24 minutes. Slides were washed in 3 changes of Discovery Wash (Roche Tissue Diagnostics, ref.

07311079001), and rinsed in running tap water for 3 minutes prior to mounting with Diamond Prolong anti-fade mountant (Thermo Fisher, ref. P36970).

A negative control slide was stained in parallel with each test tissue run, replacing the antibodies of interest with CONFIRM Negative Control Rabbit Ig (Roche Tissue Diagnostics, ref. 760-1029) and Negative Control Mouse Monoclonal Antibody (MOPC-211) (Roche Tissue Diagnostics, ref. 760-2014) as appropriate.

Stained slides were imaged at 10x magnification using the PhenoImager HT (Akoya Biosciences, version 1.0). A TMA map was applied using Phenochart (Akoya Biosciences, version 1.1.0) and single core images were acquired at 20x with the PhenoImager HT. Core images were spectrally unmixed using InForm software (Akoya Biosciences, version 2.6) using single-stained fluorescent images to generate a spectral library and an unstained autofluorescence image to remove the background.

*High-plex immunofluorescence assay*

The high-plex mIF assay was applied to a single TMA slide comprising UK cases only. Staining and imaging were performed integratively with the PhenoCycler Fusion platform (Akoya Biosciences) using the following protein markers. These markers represent a subset of a larger panel and were selected for highly focused analysis of epithelial subtypes.

SMA-BX013_RX013-AF750 (Akoya Bioscience, Ref. 4450049, clone 1A4), S100A4/FSP (Sigma-Aldrich, Ref. 07-2274, Polyclonal), IL-6 (ABCAM, Ref. 9324, Clone 1.2-2B11-2G10), LIF (Proteintech, Ref. 26757-1-AP, carrier-free version, Polyclonal), GATA6 (Cell

Signalling Technology, Ref. 5851BF, Clone D61E4), HNF4alpha (Cell Signalling Technology, Ref. 31059, Clone C11F12), CKae1/ae3-BX019_X019-AF750 (Akoya Biosciences, Ref. 4450020, clone AE-1/AE-3), FAP (Cell Signalling Technology, Ref. 81192, clone F1A4G), P40 (ABCAM, Ref. 269956, Clone EPR17863-47), S100A2 (ABCAM, Ref. 247877, Clone EPR5392), Keratin17 (Biolegend, Ref. 697202, Clone W16131A), CTGF (ABCAM, Ref. 6992, Polyclonal), FOX1A1 (ABCAM, Ref. 249749, Clone EPR10881-14), and PDGFRa, (Cell Signalling Technology, Ref. 32641, Clone D1E1E1).

All antibodies were validated in a chromogenic immunohistochemical assay (DAB) using the Ventana Discovery Ultra (RUO Discovery Ultra, Roche Tissue Diagnostics, version 21.00.0019) to determine the optimal dilution using control tissue known to express each antigen, then applied to the tissue of interest. Once validated in DAB, the antibodies were assigned to a unique PhenoCycler barcode and conjugated using the protocol below.

1. Add 500µl of Filter blocking solution (Antibody Conjugation Kit, Akoya Biosciences, Ref. 7000009) to the top of the 50kDa MWCO filter (Sigma Aldrich, Ref. UFC5050) and spin down at 12000g for 2 mins.
2. Empty both the top and the bottom of the column.
3. Add 100µg of the antibody and 300µl of PBS.
4. Spin down at 12000g for 8 mins, empty the bottom of the column.
5. Create the antibody reduction master mix:
   Reduction solution 2 (275µl) + Reduction solution 1 (6.6 µl) (Antibody Conjugation Kit, Akoya Biosciences, Ref. 7000009).
6. Add 260 µl of the reduction master mix, pipette up and down or vortex for 3 sec.
7. Incubate for 15 minutes at RT.

8. Add 200µl of conjugation solution (Antibody Conjugation Kit, Akoya Biosciences, Ref. 7000009).
9. Spin down at 12000g for 8 mins, empty the bottom of the column.
10. Add 450µl of conjugation solution, pipette up and down or vortex for 3 sec.
11. Spin down at 12000g for 8 mins, empty the bottom of the column.
12. Prepare the barcode solution:
    a. Resuspend two barcodes, each in 10µl of nuclear free water (Thermo scientific, Ref. R0582). Avoid vortex.
    b. Add 210µl of Conjugation solution to each suspended Barcode (avoid vortex, gentle pipette).
    c. Leave on the shaker.
13. Add the barcode solution to the top of the column that contains the antibody, pipette up and down or vortex for 3 sec.
14. Transfer in a low bind tube and place on the shaker, incubate overnight at 4°C.
15. Transfer the solution back in the column.
16. Spin down at 12000g for 8 mins, empty the bottom of the column.
17. Add 450µl of purification solution (Antibody Conjugation Kit, Akoya Biosciences, Ref. 7000009), pipette up and down or vortex for 3 sec.
18. Spin down at 12000g for 8 mins, empty the bottom of the column.
19. Add 450µl of purification solution, pipette up and down or vortex for 3 sec.
20. Spin down at 12000g for 8 mins, empty the bottom of the column.
21. Add 450µl of purification solution, pipette up and down or vortex for 3 sec.
22. Spin down at 12000g for 8 mins empty the bottom of the column.
23. The top of the column will contain the purified solution.

24. Add 150µl of Antibody Stabilizer PBS PROTEIN-FREE (Bioaxxess, Ref. 331), pipette up and down or vortex for 3 sec.
25. Place a new empty tube upside-down on top of the filter unit column.
26. Invert the filter for collection into the new collection tube.
27. Spin down at 3000g for 2 mins.
28. The final volume should be around 170µl.
29. Transfer in an Eppendorf.

The conjugated antibodies were tested in DAB and compared to the original DAB (with the non-conjugated antibody) for loss of signal. Each antibody was then tested in fluorescence using the PhenoCycler Fusion.

The TMA section was cut at 4µm and baked at 60°C for 1 hour. The slide was dewaxed, and antigen retrieval was performed using the Ventana Discovery Ultra (RUO Discovery Ultra, Roche Tissue Diagnostics, version 21.00.0019) with Discovery Cell Conditioning 1 (Roche Tissue Diagnostics, 06414575001) at 95°C for 32 minutes. The slide was then washed and stained using the supplied protocol (Table SM1) and with the following antibody-barcode combinations.

S100A4/FSP was assigned to BX052_RX052-ATTO550 (Akoya Biosciences, Ref. 5250012) and applied overnight at 1/25, IL-6 was assigned to BX033_RX033-AF647 (Akoya Biosciences, Ref. 5550013) and applied for 3 hours at 1/200, LIF was assigned to BX006_RX006-AF647 (Akoya Biosciences, Ref. 5550018) and applied for 3 hours at 1/200, GATA6 was assigned to BX028_RX028-AF647 (Akoya Biosciences, Ref. 555020) and applied overnight at 1/10, HNF4alpha was assigned to BX045_RX045-AF647 (Akoya

Biosciences, Ref. 5550016) and applied overnight at 1/200, FAP was assigned to BX002_RX002-ATTO550 (Akoya Biosciences, Ref. 5450023) and applied for 3 hours at 1/25, P40 was assigned to BX024_RX024-AF647 (Akoya Biosciences, Ref. 5550010) and applied for 3 hours at 1/50, S100A2 was assigned to BX040_RX040-ATTO550 (Akoya Biosciences, Ref. 5250017) ) and applied overnight at 1/100, Keratin17 was assigned to BX035_RX035-ATTO550 (Akoya Biosciences, Ref. 5250007) and applied for 3 hours at 1/100, CTGF was assigned to BX054_RX054-AF647 (Akoya Biosciences, Ref. 5550019) and applied for 3 hours at 1/300, FOX1A1 was assigned to BX020_RX020-ATTO550 (Akoya Biosciences, Ref. 5250002) and applied for 3 hours at 1/1000, PDGFRa was assigned to BX010_RX010-AF647 (Akoya Biosciences, Ref. 5550007) and applied overnight for 1/50, SMA was assigned to BX013_RX013-AF750 (Akoya Biosciences, Ref. 4450049) and applied for 3 hours at 1/300, CKae1/ae3 was assigned to BX019_RX019-AF750 (Akoya Biosciences, Ref. 4450020) and was applied for 3 hours at 1/600.

A flow cell was attached to the slide as per user guide instructions[1]. The slide was loaded in the PhenoCycler Fusion and the respective reporters were applied in cycles. A whole slide scan was acquired on the PhenoCycler Fusion imaging system in between every cycle at 20x magnification. Images were reviewed using QuPath version 0.5.0.

1. Houston, A. Akoya Biosciences PhenoCycler Fusion (formerly CODEX) User Guide v1. (2024) doi:10.17504/protocols.io.x54v9p4j1g3e/v1.

**Table SM1 | PhenoCycler Fusion staining protocol.**

| Supplier | Ref | Reagent | Incubation | Temp |
|---|---|---|---|---|
| | | dd H2O | Wash | RT |
| | | dd H2O | 10 minutes | RT |
| Akoya | 7000017 | Hydration Buffer | 2 minutes | RT |
| Akoya | 7000017 | Hydration Buffer | 2 minutes | RT |
| Akoya | 7000017 | Staining Buffer | 20 minutes | RT |
| Custom[1] | | Antibodies Cocktail 1 | 3 hours | 4ºC |
| Akoya | 700017 | Staining buffer | Wash | RT |
| Custom[1] | | Antibodies Cocktail 2 | Overnight | RT |
| Akoya | 700017 | Staining buffer | 2 minutes | RT |
| Akoya | 700017 | Staining buffer | 2 minutes | RT |
| Custom[1] | | Post staining fixing solution | 10 minutes | RT |
| | | PBS | Wash | RT |
| | | PBS | Wash | RT |
| | | PBS | Wash | RT |
| VWR | 85681.32 | Ice Methanol | 5 minutes | RT |
| | | PBS | Wash | RT |
| | | PBS | Wash | RT |
| | | PBS | Wash | RT |
| Custom[1] | | Final fixative solution | 20 minutes | RT |
| | | PBS | Wash | RT |
| | | PBS | Wash | RT |
| | | PBS | Wash | RT |
| Akoya | 700017 | Storage buffer | Up to 2 weeks | 4ºC |
